# The Glycolytic Protein Phosphofructokinase Dynamically Relocalizes into Subcellular Compartments with Liquid-like Properties *in vivo*

**DOI:** 10.1101/636449

**Authors:** SoRi Jang, Zhao Xuan, Ross C. Lagoy, Louise M. Jawerth, Ian Gonzalez, Milind Singh, Shavanie Prashad, Hee Soo Kim, Avinash Patel, Dirk R. Albrecht, Anthony A. Hyman, Daniel A. Colón-Ramos

**Author notes:** Correspondence to: Daniel A. Colón-Ramos, Ph.D., Department of Neuroscience, Department of Cell Biology, Yale University School of Medicine, 295 Congress Avenue, BCMM 436B, New Haven, CT 06510.

## Abstract

While much is known about the biochemical regulation of glycolytic enzymes, less is understood about how they are organized inside cells. Here we built a hybrid microfluidic-hydrogel device for use in *Caenorhabditis elegans* to systematically examine and quantify the dynamic subcellular localization of the rate-limiting enzyme of glycolysis, phosphofructokinase-1/PFK-1.1. We determine that endogenous PFK-1.1 localizes to distinct, tissue-specific subcellular compartments *in vivo*. In neurons, PFK-1.1 is diffusely localized in the cytosol, but capable of dynamically forming phase-separated condensates near synapses in response to energy stress from transient hypoxia. Restoring animals to normoxic conditions results in the dispersion of PFK-1.1 in the cytosol, indicating that PFK-1.1 reversibly organizes into biomolecular condensates in response to cues within the cellular environment. PFK-1.1 condensates exhibit liquid-like properties, including spheroid shapes due to surface tension, fluidity due to deformations, and fast internal molecular rearrangements. Prolonged conditions of energy stress during sustained hypoxia alter the biophysical properties of PFK-1.1 *in vivo*, affecting its viscosity and mobility within phase-separated condensates. PFK-1.1’s ability to form tetramers is critical for its capacity to form condensates *in vivo*, and heterologous self-association domain such as cryptochrome 2 (*CRY2)* is sufficient to constitutively induce the formation of PFK-1.1 condensates. PFK-1.1 condensates do not correspond to stress granules and might represent novel metabolic subcompartments. Our studies indicate that glycolytic protein PFK-1.1 can dynamically compartmentalize *in vivo* to specific subcellular compartments in response to acute energy stress via multivalency as phase-separated condensates.

## Introduction

Cells are internally organized through compartmentalization. Cellular compartments can delineate areas with specialized functions and locally concentrate molecules of a pathway (Walter and Brooks, 1995). While most of our cell biological understanding of compartmentalization is derived from the concept of membrane-bound organelles, an increasing body of work has revealed the existence of membraneless organelles—structures such as the nucleolus, germ granules, and Cajal bodies, which lack physical barriers like lipid membranes, but can still self-organize into biomolecular condensates (Banani et al., 2017; Mitrea and Kriwacki, 2016). An important question in cell biology has become which cytoplasmic processes are organized into these membraneless organelles.

Glycolysis is a fundamental energy-producing metabolic pathway that consists of ten enzymatic steps. Unlike oxidative phosphorylation, which is housed within the membrane-bound mitochondrion, glycolytic enzymes are soluble proteins in the cytosol for most eukaryotic cells. Early biochemical studies using rat muscle and brain extracts demonstrated that glycolytic proteins interchange between soluble and particulate states depending on different metabolites or hypoxic conditions, suggestive of transient interactions that could result in the subcellular organization of glycolysis (Clarke and Masters, 1974; Knull, 1978; Kurganov et al., 1985; Masters, 1984; Wilson, 1968, 1978). Studies in red blood cells, which lack mitochondria, suggest that glycolytic proteins compartmentalize near the plasma membrane and that their subcellular organization is important for the regulation of cellular cation pumps (Chu et al., 2012; Green et al., 1965; Mercer and Dunham, 1981). Immunohistochemical studies on isolated *Drosophila* flight muscle revealed a distinct pattern of co-localization of glycolytic protein isoforms to muscle sarcomeres (Sullivan et al., 2003) and vertebrate endothelial cells display enrichment of glycolytic proteins to lamellipodia and filopodia, where mitochondria are mostly absent (De Bock et al., 2013). Together, these studies and others suggest that glycolytic proteins could be compartmentalized in specific tissues (Kastritis and Gavin, 2018; Kohnhorst et al., 2017). Yet the notion of a subcellular organization for glycolytic proteins has remained controversial, largely due to the lack of studies examining the dynamic distribution of these enzymes *in vivo* (Brooks and Storey, 1991; Menard et al., 2014).

We recently reported that in *Caenorhabditis elegans* neurons, glycolytic proteins are necessary to power the synaptic vesicle cycle during energy stress (Jang et al., 2016). *In vivo* examination of the subcellular distribution of the *C. elegans* glycolytic proteins revealed that they can localize near synapses upon energy stress induced by transient hypoxia and that disruption of their subcellular localization affects their ability to power the synaptic vesicle cycle and sustain neuronal activity (Jang et al., 2016). Glycolytic enzymes were also shown to co-localize into subcellular compartments termed “G-bodies” in yeast, the formation of which was important for cellular division during conditions of persistent hypoxia (Jin et al., 2017). Similar observations in mammalian tissue culture cells also demonstrated that glycolytic enzymes co-localize into clusters (Kohnhorst et al., 2017). A human isoform of phosphofructokinase (PFK) was recently shown to assemble into tetramers that oligomerize into higher-ordered filamentous structures *in vitro* (Webb et al., 2017). In the same study, the ability for the PFK to oligomerize *in vitro* was proposed to underpin its ability to dynamically compartmentalize in tissue culture cells in response to specific metabolites. These studies in living cells indicate that glycolytic proteins are not simply diffusely distributed in the cytosol, and suggest the existence of regulatory mechanisms that organize glycolytic proteins *in vivo*. The purported organization of glycolytic proteins *in vivo* could have important consequences for the subcellular regulation of this metabolic pathway (Kastritis and Gavin, 2018; Wombacher, 1983; Zecchin et al., 2015). How proteins of the glycolytic pathway are subcellularly organized and how their localization is regulated are key questions that remain to be answered.

To examine these questions, we developed a hybrid microfluidic-hydrogel device for use with *C. elegans* to systematically examine and quantify the dynamic subcellular localization of glycolytic proteins *in vivo*. We focused our study on the rate-limiting enzyme of glycolysis, phosphofructokinase-1/PFK-1.1, and performed high-resolution *in vivo* imaging of its subcellular localization while precisely and dynamically controlling oxygen levels to transiently inhibit oxidative phosphorylation and induce acute energy stress. Using this system, we observed that in cells in which PFK-1.1 is diffusely localized in the cytosol, such as neurons, the enzyme can dynamically relocalize into biomolecular condensates in response to transient energy stress. Upon return to normoxic conditions, PFK-1.1 dispersed in the cytosol. We further determined that PFK-1.1 condensates exhibit liquid-like properties and that their molecular dynamics, including its viscosity and biophysical properties, change with prolonged hypoxic conditions. Together, our studies demonstrate that the glycolytic protein PFK-1.1 can dynamically compartmentalize *in vivo* in response to acute energy stress as phase-separated condensates.

## Results

### PFK-1.1 localizes to specific subcellular compartments *in vivo*

To better understand the subcellular localization of glycolytic proteins in living animals, we generated transgenic *C. elegans* strains expressing functional fluorophore-tagged PFK-1.1 (PFK-1.1::EGFP) under its own promoter (Figure S1A). Expression of PFK-1.1::EGFP rescues synaptic vesicle cycle defects observed in *pfk-1.1* loss of function alleles, indicating that tagged PFK-1.1 is both functional and able to recapitulate the endogenous expression pattern of the gene ((Jang et al., 2016) and data not shown). *In vivo* examination of the PFK-1.1 expression pattern revealed, as expected, broad expression of PFK-1.1 in multiple tissue types, including muscles and neurons (Figures 1A, S1B, and S1C). In most examined tissues there were high levels of cytoplasmic PFK-1.1 and no pronounced subcellular organization.

In some tissues, PFK-1.1 displayed subcellular organization. In muscles, PFK-1.1 localized into a striking pattern of longitudinal bands with alternating interdigitated foci (Figures 1B-1B’), consistent with the protein being specifically enriched at two subcellular compartments: 1) M-lines within the A band of muscle sarcomeres, and 2) dense body structures (functionally analogous to vertebrate Z-discs) (Figure 1C). M-lines and dense bodies function as anchors for thick myosin fibers and thin actin fibers, respectively, and their organization within muscle cells is necessary for filament cross-bridge formation and muscle contractions (Francis and Waterston, 1985; Gieseler et al., 2018; Moerman and Williams, 2006; Qadota and Benian, 2010). Consistent with our observations, immunohistological studies in *Drosophila* have also shown that glycolytic proteins are enriched at M-lines and Z-discs in flight muscles (Sullivan et al., 2003).

**Figure 1.**
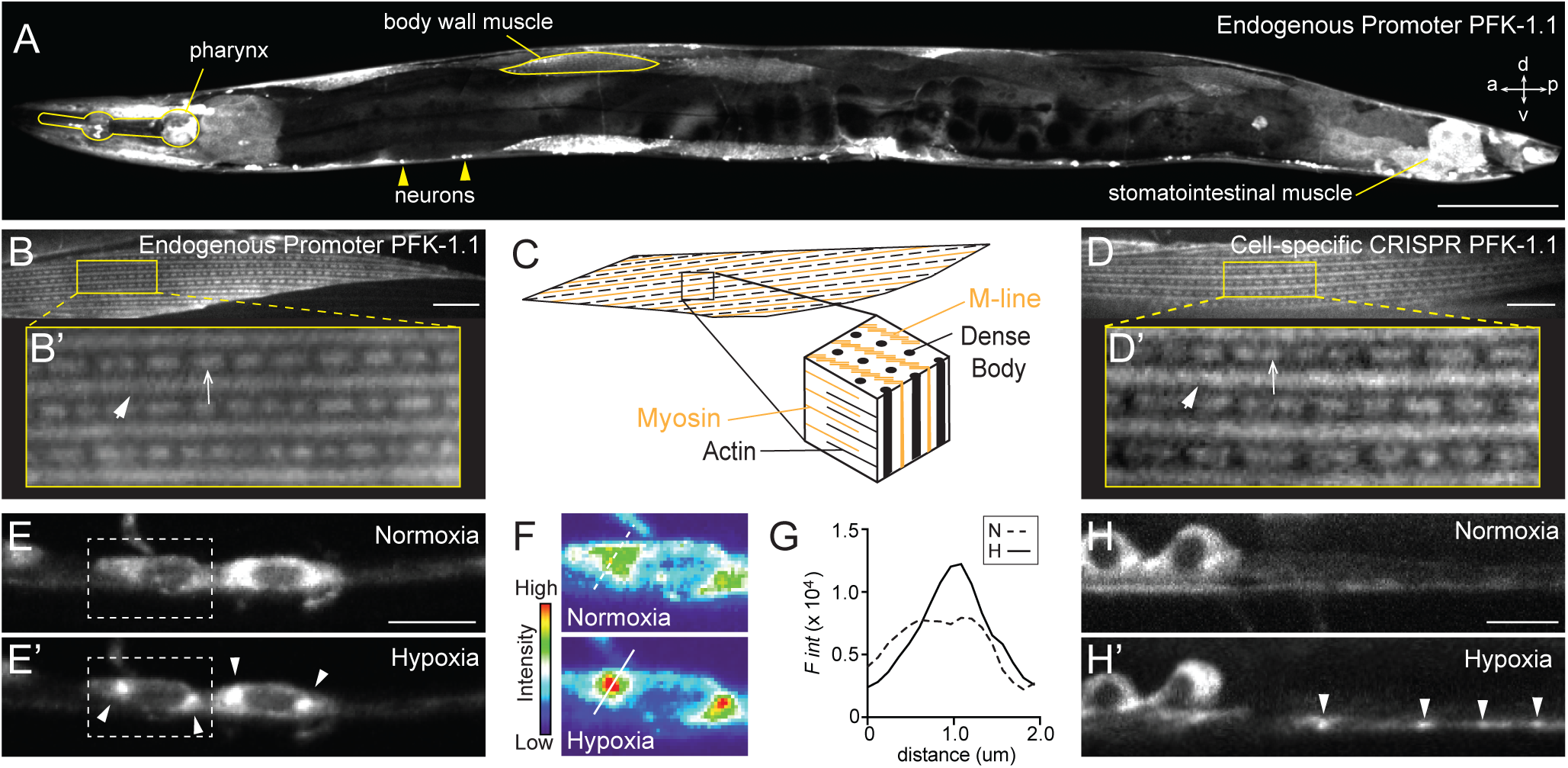
Subcellular localization of PFK-1.1 in neurons and muscles of *C. elegans*. (**A**) Localization of PFK-1.1::EGFP expressed under its own promoter in *C. elegans*, with specific tissues labeled. For orientation: anterior(a), posterior (p), dorsal (d), and ventral (v). Scale, 100 µm. (**B, B’**) *C. elegans* body wall muscle showing the localization of PFK-1.1 in M-lines (arrowhead) and dense bodies (arrow). PFK-1.1 was expressed under its own promoter. Scale, 10 µm. (**C**) Schematic showing the myofibrils of the *C. elegans* body wall muscle. M-line and dense body acts as anchors for myosin and actin filament, respectively. (**D-H’**) The *mig-13* promoter was used in conditional CRISPR lines to tag the endogenous PFK-1.1 with GFP in a subset of tissues. (**D, D’**) Endogenous PFK-1.1 also localizes to M-lines (arrowhead) and dense bodies (arrow) in the body wall muscle. Scale, 10 µm. (**E, E’**) PFK-1.1 clusters form in the neuronal cell bodies (arrowheads) over time under hypoxia. Here, elapsed time of hypoxia was 55 minutes. (**F**) Pseudo-color based on fluorescence intensity of the dashed box in **E** for the two conditions. (**G**) Quantification of fluorescence intensity along the lines shown in **F** for each condition: N (normoxia) and H (hypoxia). (**H, H’**) PFK-1.1 clusters can be seen in the neurites (arrowheads) after hypoxia. Here, elapsed time of hypoxia was 10 minutes.

To better examine the *in vivo* subcellular localization of endogenous PFK-1.1 in single cells, we generated conditional transgenic lines via CRISPR-Cas strategies (Dickinson et al., 2015; Flavell et al., 2013; Schwartz and Jorgensen, 2016). In brief, we used CRISPR-Cas to introduce two flippase recombinase target (FRT) sites that flank a transcriptional stop motif followed by a GFP sequence to the C-terminus of the endogenous *pfk-1.1* locus (Figure S1D). Introduction of these FRT-sites did not result in detectable mutant phenotypes, including the synaptic vesicle phenotype observed for the PFK-1.1 loss of function alleles (data not shown), suggesting that the genomic modifications do not alter PFK-1.1 expression or function. Using cell-specific promoters, we then drove the expression of flippase (FLPase) to excise the FRT-flanking transcriptional stop motif, allowing for the expression of the endogenous GFP-tagged PFK-1.1 in single cells (see Methods). FLPase expression using *mig-13* and *unc-47* promoters resulted in the expression of PFK-1.1::GFP in subsets of tissues, and with a low signal due to endogenous labeling. Consistent with the localization pattern of the overexpressed PFK-1.1::EGFP (Figure 1B), we also observed the localization of PFK-1.1 at M-lines and dense bodies in the muscles of the cell-specific CRISPR lines (Figures 1D-1D’).

We then examined the subcellular localization of PFK-1.1 in neurons. Expression of PFK-1.1:EGFP in single neurons revealed that, while PFK-1.1 is not equally distributed throughout the neurite, it is for the most part diffusely localized in the cytosol (Figure S1E). Previously we observed that under conditions known to cause energy stress, such as transient hypoxia, PFK-1.1 dynamically relocalizes to subcellular compartments (Jang et al., 2016). These former studies were done by over-expressing PFK-1.1 from cell-specific promoters in single neurons. Consistent with these observations, an examination of the conditional CRISPR lines revealed that, upon transient hypoxia, endogenous PFK-1.1 relocalizes from a diffuse cytosolic pattern to subcellular clusters both in the neuronal soma and in neurites (Figures 1E-1H’ and S1F).

Our findings demonstrate that *in vivo* and in metazoans, PFK-1.1 localizes to specific subcellular compartments. They also reveal that the localization pattern of PFK-1.1 varies between tissues. Importantly, our studies demonstrate that even in cells that have no obvious subcellular organization of PFK-1.1, such as the neurons, the localization of endogenous PFK-1.1 can dynamically change in response to cues within the cellular environment.

### PFK-1.1 dynamically relocalizes to distinct subcellular compartments in response to transient hypoxia

Transient hypoxia inhibits oxidative phosphorylation at the mitochondria and induces cellular energy stress. To characterize the dynamic responses of PFK-1.1 to controlled changes in oxygen concentration, we developed a hybrid microfluidic-hydrogel device that enables precise regulation of oxygen concentration in the animal’s surrounding environment during high-resolution and long-term imaging of PFK-1.1 localization (Figures 2A-2C). Calibration of our device enabled us to make precise and rapid switches between steady-state hypoxic (0% O_2_) and normoxic (21% O_2_) conditions within 1 minute (Figures S2A-S2D). Animals under sustained steady-state normoxic conditions (21% O_2_) did not show relocalization of PFK-1.1 proteins in neurons (Figure S3A), indicating that variables such as air flow or mechanical forces during mounting to the device do not affect the localization of PFK-1.1. Induction of transient hypoxia, however, resulted in robust relocalization of PFK-1.1 in neurons, from a diffuse state in the cytoplasm into distinct clusters near synapses (Figures 2D). The increased signal of PFK-1.1 near synapses was not the result of an increase in total protein levels at the neurite, as the total fluorescence in the neurite did not increase upon hypoxic stimuli (Figure S2E).

Cellular stress is known to cause the aggregation of cytoplasmic RNAs and proteins into stress granules (Protter and Parker, 2016). To examine if PFK-1.1 in neurons was relocalizing into stress granules, we simultaneously imaged PFK-1.1 and the stress granule protein, TIAR-1, under heat shock conditions known to cause stress granule formation (Huelgas-Morales et al., 2016; Sun et al., 2011). Consistent with previous studies, TIAR-1 in neurons was enriched in the soma and could be seen in both the cytoplasm and nuclei (Figure 2E). Heat shock, by incubating worms at 37°C for 1 hour, resulted in the aggregation of TIAR-1 in both the cytoplasm and nuclei, as expected. However, PFK-1.1 did not form clusters under conditions that caused TIAR-1 aggregation (Figure 2F). We then examined whether stress granules are formed upon transient hypoxic conditions known to induce PFK-1.1 clustering. We observed that while transient hypoxia caused PFK-1.1 clustering, the same conditions did not cause TIAR-1 clustering (Figure 2G). Our findings demonstrate that TIAR-1 and PFK-1.1 cluster under different cellular conditions, and suggest that PFK-1.1 clusters correspond to a new subcellular compartment distinct from stress granules.

**Figure 2.**
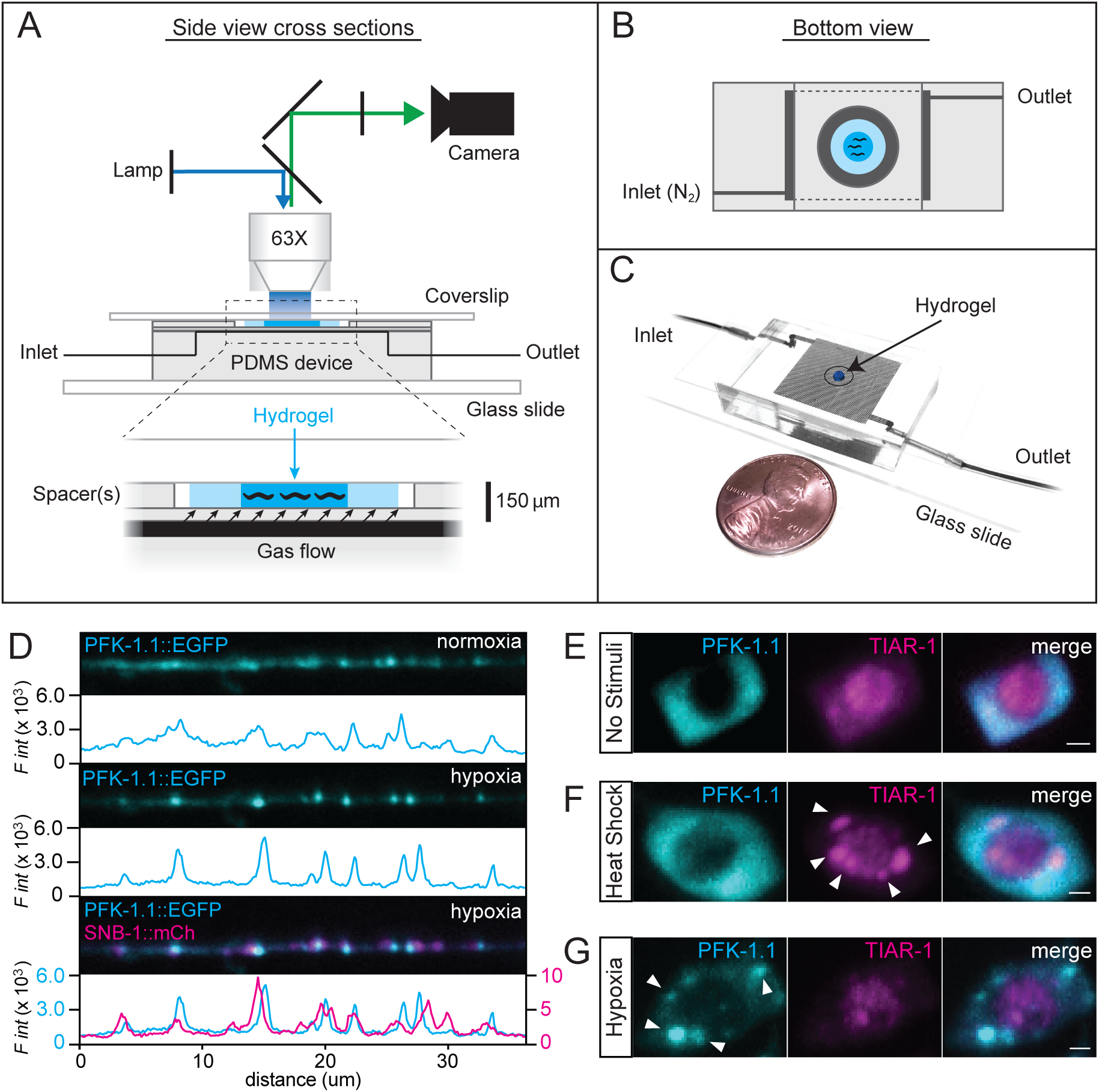
Hybrid microfluidic-hydrogel device enables high-resolution imaging of PFK-1.1 localization and clustering dynamics during transient hypoxia. (**A**) Schematic of the hybrid microfluidic-hydrogel device used to induce transient hypoxic conditions in *C. elegans* while imaging PFK-1.1 continuously and at high-resolution. Zoom-in (below) shows micron-scale device geometries for animal immobilization, gas delivery, and imaging. (**B**) A bottom view schematic of the device shows geometry of inlet and outlet channels connected to the gas arena (dark gray area). Animals are immobilized in hydrogel (dark blue) surrounded by buffer (light blue) and sealed by a circular, hole-punched PDMS spacer. Oxygen conditions experienced by the animals are controlled by inlet gas flow. (**C**) A picture of the microfluidic device, size comparable to a penny, with inlet and outlet tubing connected. (**D**) PFK-1.1::EGFP (cyan) in GABAergic neuron under normoxia (top panels) and after 10 minutes of transient hypoxia (middle panels). Co-expression with synaptobrevin-1/SNB-1::mCh (magenta, lower panel) shows that PFK-1.1 relocalized into clusters near synaptic sites upon transient hypoxia. Corresponding fluorescence intensity for each image is shown immediately below the corresponding image, with colors matching traced protein distribution in the image. (**E**) PFK-1.1::mRuby-3 (cyan) and TIAR-1::EGFP (magenta) in the cell body of neurons with no stimuli (normoxic conditions and 22°C). (**F**) Heat shock (37°C for 1 hour) leads to formation of stress granules as observed with TIAR-1 aggregates (arrowheads). PFK-1.1 does not form clusters under these condition. (**G**) Under transient hypoxic conditions, PFK-1.1 forms clusters in the cell body (arrowheads) that do not co-localize with TIAR-1. All scales, 1 µm.

### PFK-1.1 reversibly organizes into biomolecular condensates

To examine the dynamics of PFK-1.1 clusters, we performed high-resolution time-lapse imaging of PFK-1.1 in single neurons of living animals upon transient hypoxia exposure. We focused our analyses in 3-micron neurite regions that displayed PFK-1.1 enrichment during normoxia. We observed that induction of transient hypoxia resulted in a redistribution of the PFK-1.1 protein within these regions, with PFK-1.1 concentrating into 0.5-1 micron compartments (Figures 3A-3B). We also observed a concomitant decrease of PFK-1.1 levels in regions adjacent to the emerging compartments (Figure 3C). Kymograph of PFK-1.1 in neurites showed that the local formation of PFK-1.1 clusters, which remained relatively immobile, was due to a redistribution of diffuse PFK-1.1 proteins to concentrated compartments (Figure 3D). Together, our results indicate that PFK-1.1 puncta emerge through the local condensation of PFK-1.1 material into concentrated clusters, which we now refer as condensates (Banani et al., 2017; Shin and Brangwynne, 2017).

**Figure 3.**
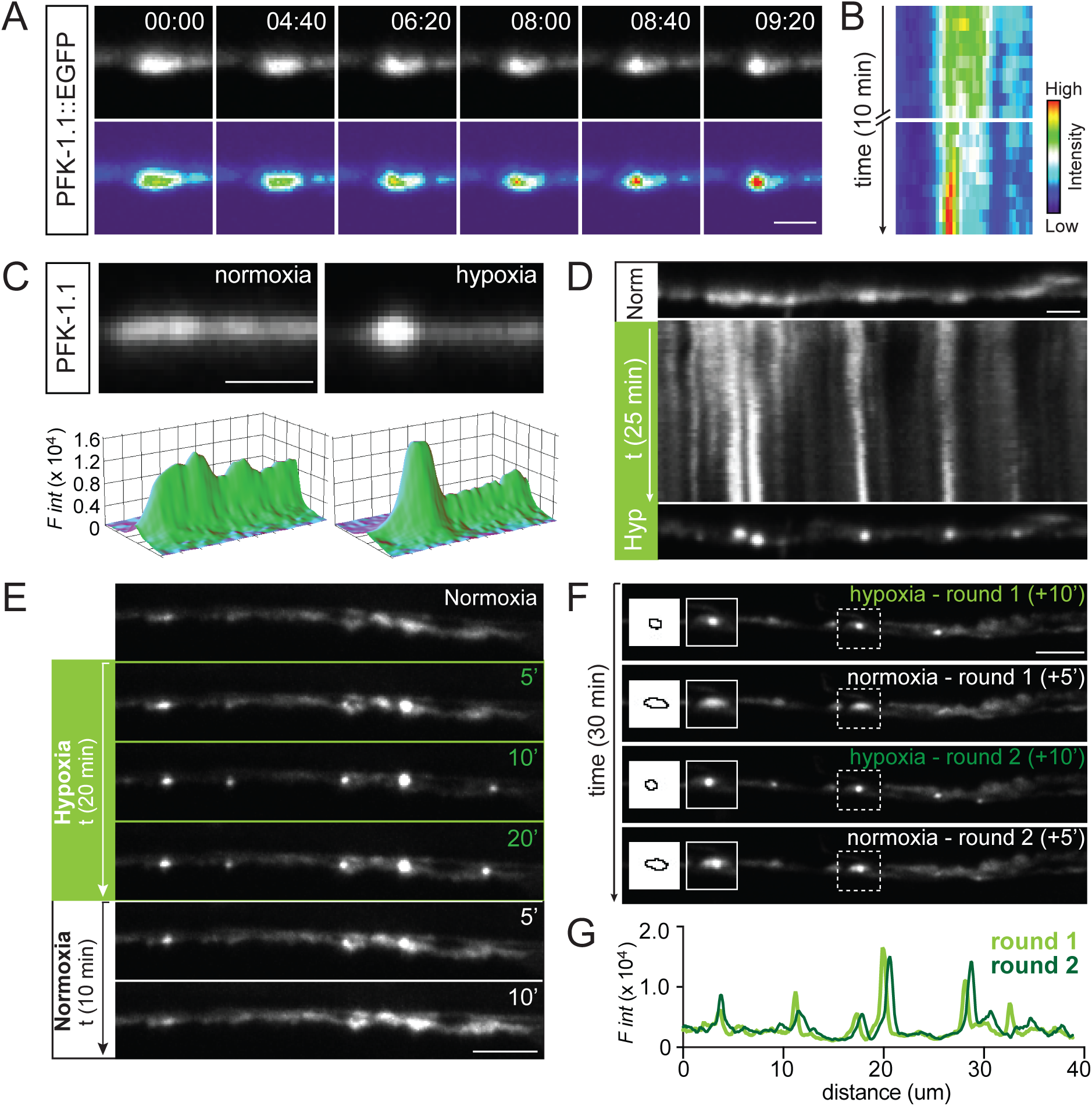
PFK-1.1 reversibly organizes into biomolecular condensates. (**A**) Upon minutes of transient hypoxia, PFK-1.1 relocalizes from a dispersed localization pattern in the cytosol to a concentrated punctum. Time elapsed is indicated for each image in minutes:seconds. Bottom images have been pseudo-colored based on fluorescence intensity. Scale, 2 µm. (**B**) Kymograph of PFK-1.1 localization shown in **A** over 10 minutes of transient hypoxia. 0-3 minutes and 7-10 minutes are shown. (**C**) Localization of PFK-1.1 in a neurite during normoxia (left panel) and after 18 minutes of transient hypoxia (right panel). Corresponding 3-D surface rendering projections of fluorescence intensity in panels below. Scale, 1 µm. (**D**) Kymograph showing PFK-1.1 localization change over time, starting from normoxic condition (Norm, top panel) to 25 minutes of transient hypoxia (Hyp) (kymograph and lower panel). Scale, 2 µm. (**E**) Localization of PFK-1.1 through a normoxia-hypoxia-normoxia cycle at different time points (as indicated by minutes in upper right hand of images). PFK-1.1 condensates induced by transient hypoxic condition (green numbers, in minutes) can be seen dispersing away when the condition is returned to normoxia (white numbers, in minutes). See also Movie 1. Scale, 5 µm. (**F**) PFK-1.1 localization under two cycles of transient hypoxia (round 1 and 2). PFK-1.1 forms condensates during 10 minutes of transient hypoxia and becomes diffuse during 5 minutes of normoxia. Dashed box in main panel corresponds to zoomed in image on the left (along with the outline of the corresponding morphology of the condensate). See also Movie 2. Scale, 5 µm. (**G**) Line scan of first round (light green) and second round (dark green) of PFK-1.1 condensate localization in the neurite shown in **F** under transient hypoxic conditions. Note that during repeated cycles of normoxia and transient hypoxia, PFK-1.1 condensates reappear at similar subcellular locations.

We observed that condensates of PFK-1.1 were visible within five minutes of transient hypoxia. The persistence of hypoxic treatment resulted in the emergence of additional condensates, resulting in the almost complete relocalization of all visible PFK-1.1::EGFP into clusters (Figure 3E). Interestingly, restoring animals to normoxic conditions resulted in the dispersion of PFK-1.1 in the cytosol in both the neurite and cell body (Figure 3E, Movie 1 and Figures S3B). Thus, we asked whether PFK-1.1 proteins were capable of repeated cycles of condensation and dispersion in neurons. We observed that PFK-1.1 could undergo multiple cycles of condensation, displaying similar dynamics in each cycle (Figure 3F and Movie 2). Importantly, during the two rounds of transient hypoxia and normoxia, we observed that PFK-1.1 condensates reappeared in similar locations (Figures 3G and S3C). The repeated formation of PFK-1.1 clusters at the same location in the neurite occurred even after PFK-1.1 proteins completely diffused back to their initial state (Figure S3D). The reappearance of PFK-1.1 clusters at the same sites is consistent with our observations that PFK-1.1 clusters form preferentially (but not exclusively) near synapses ((Jang et al., 2016) and Figure 2D). The observation may suggest the existence of synaptic nucleating factors, or the existence of subcellular conditions at the synapse which instructs local assembly of PFK-1.1 upon transient hypoxia (Berry et al., 2018).

### PFK-1.1 condensates exhibit liquid-like behaviors

Biomolecular condensates, including the nucleolus, germ granules, prion proteins, and postsynaptic proteins, have been shown to exhibit liquid-like characteristics (Brangwynne et al., 2009, 2011; Franzmann et al., 2018; Patel et al., 2015; Zeng et al., 2016). To examine if PFK-1.1 condensates transiently induced by hypoxic treatment also exhibit liquid-like properties, we tested for the three key features that define liquid-like compartments: 1) Liquid-like compartments are spherical due to surface tension, 2) they can fuse and relax into a single spherical droplet, and 3) they exhibit fast internal molecular rearrangements (Alberti et al., 2019). We systematically tested the PFK-1.1 clusters for their biophysical characteristics.

#### 1) Liquid-like compartments are spherical due to surface tension

Our qualitative observations on the emerging clusters revealed that most of the PFK-1.1 condensates were circular (and, since there was no obvious *z*-axis asymmetry, likely spherical). To examine the shape of the PFK-1.1 condensates, and to establish a quantitative criterion that defines the emergence of the clusters during condensation of PFK-1.1, we computed the aspect ratio, a ratio between the major and minor axes of the emerging clusters, over time (Figures 4A, S4A, and S4B) (Brangwynne et al., 2011; Gopal et al., 2017). Aspect ratio calculations of hypoxia-induced PFK-1.1 clusters in neurites revealed a distribution with a mean aspect ratio of 1.32 +/- 0.04 (+/- SEM; N=38) (Figure 4B). A perfect sphere has an aspect ratio of 1, so our data suggest that PFK-1.1 clusters were similar, on average, to elongated spheroids.

**Figure 4.**
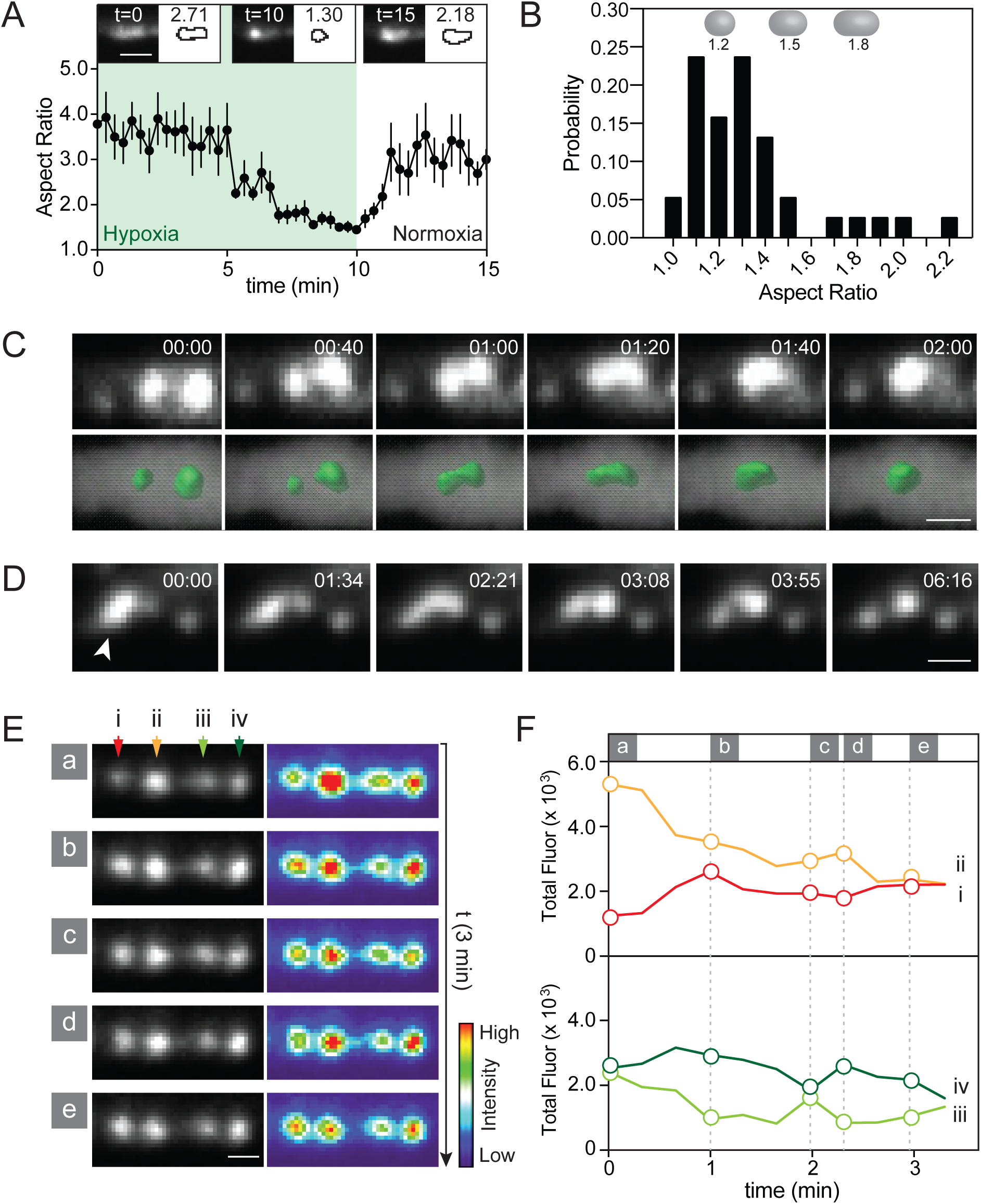
PFK-1.1 condensates exhibit liquid-like behaviors. (**A**) Reversibility of PFK-1.1 localization as quantified by the aspect ratio of a population of PFK-1.1 condensates through one cycle of hypoxia-normoxia (N=8 condensates). Mean (black circles) and SEM values are shown for each time point. Images on top, and corresponding outlines, are representative morphologies of condensates, from left to right, the beginning of hypoxia, after 10 minutes of transient hypoxia, and after 5 minutes of recovery in normoxia. Scale, 2 µm. (**B**) Histogram of the probability distribution of the PFK-1.1 condensate aspect ratio after 40 minutes of transient hypoxia (N=38 condensates). A perfect sphere has an aspect ratio of 1, and PFK-1.1 condensates have an average aspect ratio of 1.3. Representative spherocylindrical shapes for different aspect ratios (1.2, 1.5, and 1.8) are shown above. (**C**) Time-lapse images of adjacent PFK-1.1 condensates under transient hypoxic conditions fusing and relaxing into a spheroid (above) and their corresponding 3-D surfaced rendered images (below). Elapsed-time from the beginning of the imaging session is shown for each image in minutes:seconds. Here, 00:00 is 08:40 into the hypoxic treatment. See also Movie 3. Scale, 1 µm. (**D**) A fluid-like movement recorded for a PFK-1.1 condensate (arrowhead) during transient hypoxia. Elapsed time from the beginning of the imaging session is indicated for each image in minutes:seconds. Scale, 1 µm. (**E**) Four adjacent PFK-1.1 condensates (i-iv) during transient hypoxia displayed exchange of PFK-1.1::EGFP in the span of three minutes. The first image (a) is after approximately six minutes into hypoxic treatment. See also Movie 4. Scale, 1 µm. (**F**) Quantification of the fluorescence of the PFK-1.1 condensates in **E**, with graphs pseudocolored to corresponding arrowheads in (i-iv) and timepoints (a-e) in **E**.

Because we qualitatively also observed a relationship between the size of the puncta and the extent of the elongated spheroids (Figure S4C), we hypothesized that neurite space constraints could affect the shape of the larger (liquid) PFK-1.1 condensates. Liquids are capable of deformation, and liquid droplets confined within a cylindrical-like space, such as a neurite, would be expected to elongate into capsule-shaped spherocylinders. To test if the larger PFK-1.1 condensates were constrained capsule-shaped spherocylinders, we first calculated the average diameter of the neurites by measuring cytoplasmic mCherry expressed in GABAergic neurons, and by examining electron micrographs of GABAergic neurites (Xuan et al., 2017). From these calculations, we determined the average diameter of the examined neurites to be 0.64 +/- 0.06 µm (mean +/- SEM; N=20) (Figures S4D). Empirical measurement of the area of the PFK-1.1 condensates (see Methods) revealed that these structures had an average area of 0.43 +/- 0.04 µm^2^ (mean +/- SEM; N=38) (Figure S4E). Assuming radial symmetry of the neurite, average condensate volume would be 0.21 +/- 0.03 µm^3^, which is larger than the maximum sphere fitting within the neurite diameter, 0.14 +/- 0.04 µm^3^. A hypothetical spherocylinder with a volume of 0.21 µm^3^ and radial diameter of 0.64 µm would have a length of 0.87 µm and an aspect ratio of 1.36 (Figure S4F), a value similar to the one we empirically calculated, which was 1.32. Thus, our findings suggest that PFK-1.1 condensates have liquid-like properties that result in spherocylindrical morphologies due to constraints by two physical forces: the surface tension of the droplet and the dimensions of the neurite.

#### 2) Liquid-like structures can fuse and relax into a single spherical droplet

Phase separated compartments display stereotypical fluid-like behaviors that include fusion. We reasoned that if PFK-1.1 condensates displayed liquid-like characteristics, condensates that happened to nucleate in close proximity (within one micron from each other) would fuse. Indeed, PFK-1.1 condensates that came into physical contact fused into new, larger clusters (Figures 4C and S5A, and Movies 3 and 4). The new clusters relaxed into spheroid shapes upon fusion, consistent with a thermodynamically-favored response of a liquid compartment to surface tension (Brangwynne et al., 2009). Moreover, while most PFK-1.1 particles *in vivo* retained spheroid shapes upon formation, some particles would occasionally become deformed and even shear, presumably because of cytoplasmic flux or transport of organelles disrupting the PFK-1.1 condensates in the tightly-packed spaces of the neurite (Figure 4D). Interestingly, deformation of the condensates resulted in transient fluid-like movements and the eventual relaxation back into spheroid structures.

Quantification of the levels of fluorescence in nearby condensates revealed that growth of a condensate happened at the expense of adjacent condensates in a process that was reminiscent of Ostwald ripening (Figure 4E). But unlike Ostwald ripening, which is a thermodynamically spontaneous process where smaller droplets dissolve into energetically-favored larger droplets (Voorhees, 1992), some PFK-1.1 condensates *in vivo* did not result in larger droplets. Instead, the exchange of material reached a dynamic equilibrium between adjacent condensates as they approached similar fluorescence levels (Figures 4E, 4F, and Movie 5). The observed *in vivo* dynamics between adjacent condensates could be because of occlusion of puncta in the tube-like geometry of the neurite from active cellular processes that drive the formation of two similar condensates (Berry et al., 2018; Weber et al., 2019), or because of other unidentified phenomena. Importantly, the exchange of material between adjacent clusters is consistent with liquid-like behaviors of the PFK-1.1 condensates.

#### 3) Liquid compartments exhibit fast internal molecular rearrangements

To further probe the molecular dynamics of PFK-1.1 condensates, we performed fluorescence recovery after photobleaching (FRAP) experiments. We observed that upon partial FRAP of the PFK-1.1 condensate, fluorescence in the bleached area recovered within seconds of post-bleaching (Figures 5A-5B and S5B-S5B’). Consistent with an internal rearrangement resulting in fluorescence recovery, we observed that fluorescence in the bleached region increased at the expense of the unbleached areas (Figure 5C). The observed molecular dynamics for PFK-1.1 within condensates suggest that PFK-1.1 remains in liquid phase droplets capable of undergoing internal dynamic rearrangements (Brangwynne et al., 2009).

**Figure 5.**
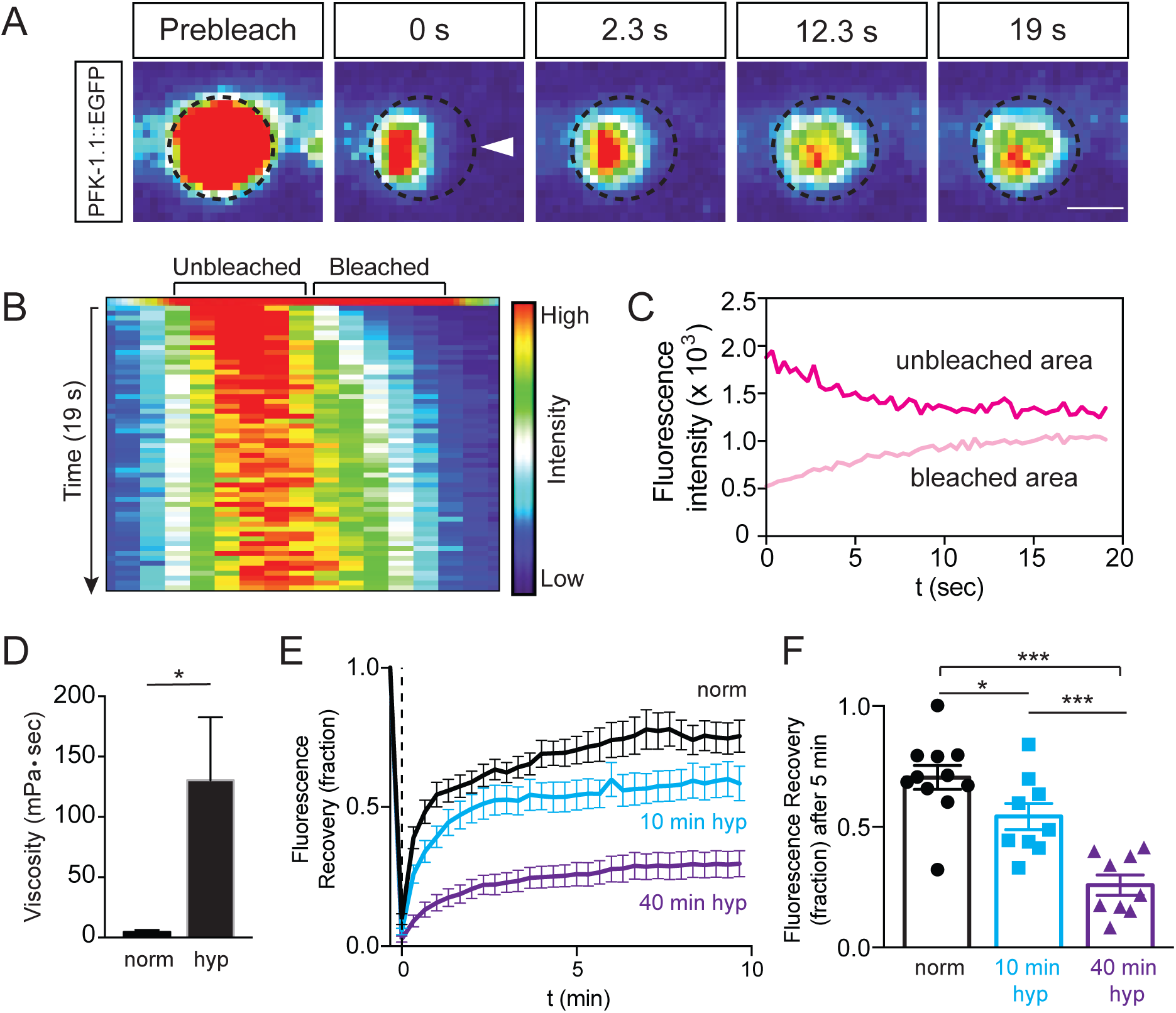
PFK-1.1 compartments exhibit fast internal molecular rearrangements that harden with time. (**A**) Fluorescence recovery after photobleaching (FRAP) of a PFK-1.1::EGFP condensate after 10 to 20 minutes of transient hypoxia. PFK-1.1::EGFP condensate initial shape outlined with dashed circle, and partial area bleached highlighted with arrowhead in second panel. Scale, 1 µm. (**B**) Kymograph of fluorescence distribution across the bleached punctum in **A**. (**C**) Change in fluorescence values for the unbleached region (dark pink line) and the bleached region (light pink line) of the condensate in **A** over time (seconds). (**D**) Viscosity estimates of PFK-1.1 proteins under normoxia (labeled “norm”) and 10 to 20 minutes of transient hypoxia (labeled “hyp”). (N=4 neurons for each conditions). (**E-F**) Fluorescence recovery after photobleaching of cytosolic PFK-1.1 after normoxia (black line, labeled “norm”), PFK-1.1 condensate after 10 minutes of transient hypoxia (blue line, labeled “10 min hyp”), and PFK-1.1 condensate after 40 minutes of transient hypoxia (purple line, labeled “40 min hyp”), and corresponding fraction of fluorescence recovery 5 minutes post-bleaching calculated in **F**. In all experiments, PFK-1.1 was photobleached to a minimum of 80% of its original fluorescence value. Note how recovery dynamics decrease with increased exposure to transient hypoxia, suggesting that PFK-1.1 condensates harden with time. Error bars denote SEM. *, p < 0.05. **, p < 0.01. ***, p < 0.001 between indicated groups.

### PFK-1.1 condensates harden with time

To test if transient hypoxia was inducing a change in the molecular dynamics of PFK-1.1 resulting in liquid-liquid phase separation, we measured the half-time recovery and lateral diffusion of PFK-1.1 in its soluble state during normoxia versus its condensate state during hypoxia (Brangwynne et al., 2009). Using FRAP we determined that the average half-time recovery in the photo-bleached regions for diffusely localized PFK-1.1 was 0.6 second while in the condensate it was 3.8 seconds (data not shown). Our findings indicate a substantial change in the diffusion dynamic of PFK-1.1 between normoxia and hypoxia, and are consistent with changes in molecular dynamics during condensation.

To approximate the fluidity of PFK-1.1 under the two states, we used the Stokes-Einstein equation (see Methods) to estimate the viscosity of PFK-1.1. While the accuracy of the viscosity calculations is limited by the small size of the puncta, we note that, in the soluble state, PFK-1.1 had an estimated viscosity of 4.8 +/- 1.7 mPa•s (mean +/- SEM; N=4), which is approximately 5 times more viscous than water. This value is comparable to the reported viscosity (2.0-3.0 mPa•s) of the cytoplasm (Mastro et al., 1984), consistent with PFK-1.1 being largely diffusely localized in the cytosol under normoxic conditions. Upon transient hypoxic conditions, however, the calculated viscosity for PFK-1.1 was 130.5 +/- 52.1 mPa•s (mean +/- SEM; N=4) (Figure 5D). Our findings suggest that PFK-1.1 increases two orders of magnitude in viscosity after it forms condensates. Importantly, our findings indicate that the biophysical properties of PFK-1.1 change upon transient hypoxia, and suggest that these changes could contribute to PFK-1.1 relocalizing into phase-separated condensates.

Changes in the viscoelastic proteins of molecular condensates contribute to liquid-liquid phase separation. Yet it has also been documented that changes in the material phase of the condensates can contribute to their maturation into more “gel-like” or “solid-like” states that eventually result in the (pathological) aggregation of proteins (Feric et al., 2016; Franzmann et al., 2018). We reasoned that if transient hypoxia was inducing a change in the molecular dynamics of PFK-1.1, one would expect that conditions of prolonged hypoxia would further alter the molecular dynamics and the biophysical properties of the condensates. To test if prolonged hypoxia alters the PFK-1.1 dynamics, we compared the fluorescence recovery of PFK-1.1 condensates that were photobleached after 10 minutes (newly formed condensates) and 40 minutes (“mature” condensates) of hypoxic treatment. We observed that the recovery dynamics of mature PFK-1.1 condensates were significantly slower than those of newly formed condensates (Figures 5E-5F).

In the course of our studies, we also imaged mature condensates which happened to be adjacent to one another. These condensates sometimes came onto physical contact with each other, just as we had previously documented for the fusion events occurring in newly formed condensates (Figure 4C). Interestingly however, and unlike the newly formed condensates, some of these mature condensates deformed and “bounced” off each other upon contact (Figure S5C and Movie 6). Our observations of these dynamics are consistent with the viscosity analyses and suggest that the viscoelastic properties of PFK-1.1 condensates change upon persistent hypoxia.

### Local concentration of PFK-1.1 drives the initiation of condensates

Physical principles of phase transitions influence the assembly of membraneless organelles such as p-granules and the nucleolus (Hyman et al., 2014). A key biophysical aspect that influence the dynamics of membraneless organelle assembly is protein concentration (Alberti et al., 2019; Banani et al., 2016; Shin and Brangwynne, 2017). Because we observed a nonhomogeneous distribution of PFK-1.1 proteins throughout the neurite even during normoxia (Figure S1E), we hypothesized that PFK-1.1 concentration may influence cluster formation at subcellular regions. To test this hypothesis, we monitored the formation of PFK-1.1 condensates for specific subcellular regions that had varying concentrations of PFK-1.1 under conditions of persistent hypoxia (see Methods). From the lengthened hypoxic treatment, we observed a continuous and asynchronous formation of PFK-1.1 condensates throughout the neurite, with PFK-1.1 condensates forming at different time points in different cellular regions (Figures 6A-6B and Movie 7). Comparisons of the local concentration of PFK-1.1 proteins in specific subcellular regions of the neurite, and the time at which the PFK-1.1 condensates initiate in those regions, revealed an inverse correlation between these two variables (Figure 6C). Our findings are consistent with initial PFK-1.1 concentrations affecting the time of initiation of the condensates, with a higher level of PFK-1.1 proteins corresponding to an earlier appearance of PFK-1.1 condensates.

**Figure 6.**
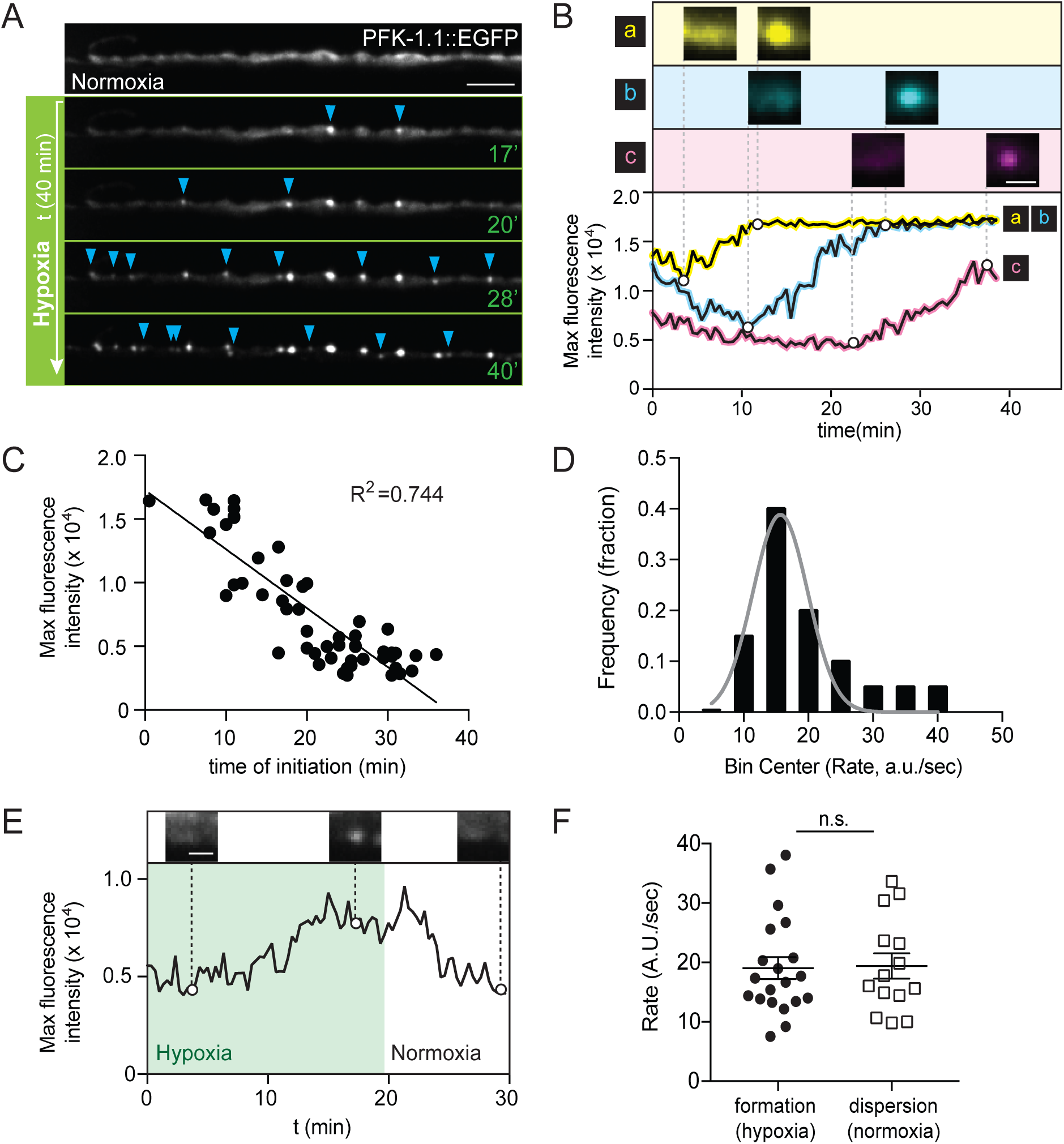
Local concentration of PFK-1.1 drives initiation of condensates. (**A**) PFK-1.1 dynamics through persistent hypoxia. Blue arrowheads mark newly formed PFK-1.1 condensates at each time point (green numbers indicate minutes into hypoxic treatment), indicating asynchronous formation of PFK-1.1 condensates throughout the neurite. Scale, 5 µm. (**B**) Three separate regions in neurites, where PFK-1.1 condensates appear, are shown (a, b, and c). The corresponding maximum fluorescence value over time for the three regions are shown below, pseudocolored to correspond to the region specified above (also labeled with a, b and c). PFK-1.1 condensate formation occurred with a linear increase in maximum fluorescence intensity through time in response to transient hypoxia. Scale, 1 µm. (**C**) The maximum fluorescence intensity of PFK-1.1::EGFP just before PFK-1.1 condensates are observed were plotted and compared to the time when PFK-1.1 condensates first appear (time of initiation). Note how the time of initiation negatively correlates with the maximum fluorescence intensity (equivalent to the amount of PFK-1.1 in a specific region of the neurite where PFK-1.1 condensates appear). (**D**) Rates for PFK-1.1 condensate formation were calculated for 20 condensates as maximum fluorescence intensity change over time, and their distribution plotted. The rates are constant, regardless of initial concentration, and a normal (Gaussian) distribution (N=20 condensates). (**E**) Maximum fluorescence intensity of a PFK-1.1 condensate calculated over time (minutes) during transient hypoxia (shaded green) and subsequent return to normoxic conditions. Representative images of PFK-1.1 condensates shown for given timepoints above the graph. (**F**) Rate of formation and dispersion of PFK-1.1 condensates under transient hypoxia (green circles, left) and normoxia (white squares, right), respectively. Note how the rates of formation and dispersion are both constant, normally distributed (compare to **D**) and not significantly (specified as n.s. in graph) different.

Interestingly, we also observed that once PFK-1.1 condensates appeared, and regardless of where or when they appeared in the neurite, they underwent similar linear rates of increase in protein fluorescence. The constant increase of fluorescence did not vary based on the starting concentration and instead displayed a normal distribution around an average constant rate (Figures 6D). Furthermore, calculations of the rates of PFK-1.1 dispersion upon return to normoxic conditions also showed a normal distribution around an average rate that was similar to the average rate of condensation (Figures 6E-6F).

Together, we found two properties regarding the local emergence of PFK-1.1 condensates across subcellular regions: 1) concentration, which correlates with, and likely determines, the timing of initiation of PFK-1.1 condensates and 2) rates of PFK-1.1 condensate growth (during transient hypoxia) and dispersion (during normoxia), which share a similar normal distribution regardless of initial concentration at subcellular locations. Thus, the asynchronous formation of PFK-1.1 condensates throughout the neurite likely results from the initial uneven distribution of PFK-1.1 in neurites, which in turn determines the local concentration and the initiation time of PFK-1.1 condensates.

### Multivalent interactions underlie PFK-1.1 condensate formation

Multivalent interactions between proteins, or proteins and RNAs, play important roles in driving condensates into liquid-like protein droplets (Banani et al., 2017). Multivalent interactions can be mediated via folded protein domains, or intrinsically disordered regions (IDRs) with multiple interacting motifs (Alberti et al., 2019; Elbaum-Garfinkle et al., 2015; Langdon et al., 2018; Li et al., 2012; Nott et al., 2015; Smith et al., 2016; Wang et al., 2018; Zhang et al., 2015). These interactions leading to phase separation can be recapitulated *in vivo* and *in vitro* by engineering self-association domains (Bracha et al., 2018; Li et al., 2012; Shin et al., 2017), underscoring the importance (and sufficiency) of multivalent interactions in the formation of condensates. To understand what molecular driving force leads to the biophysical changes observed in PFK-1.1 condensates, we examined whether multivalent interactions of PFK-1.1 play a role in the formation of PFK-1.1 condensates.

*C. elegans* PFK-1.1 does not possess canonical domains previously associated with phase separation, such as intrinsically disordered regions, RNA binding domains or SRC homology (SH3/SH2) domains (data not shown and (Banani et al., 2017)). Instead, PFK forms tetramers through hydrophobic interfaces that facilitate multivalent interactions (Webb et al., 2015). Furthermore, PFK tetramers have dihedral symmetry and can associate into higher-order oligomeric structures (Garcia-Seisdedos et al., 2017; Sola-Penna et al., 2010; Webb et al., 2017). In the human PFK Platelet isoform, the conserved residue F649 is essential for hydrophobic interactions leading to tetramer formation, and disruption of this residue abrogates PFK biochemical activity *in vitro* (Webb et al., 2015). The equivalent residue in *C. elegans*, F674, (Figures 7A and S6A) had not been tested for its role in the oligomerization of PFK-1.1 or for its function.

**Figure 7.**
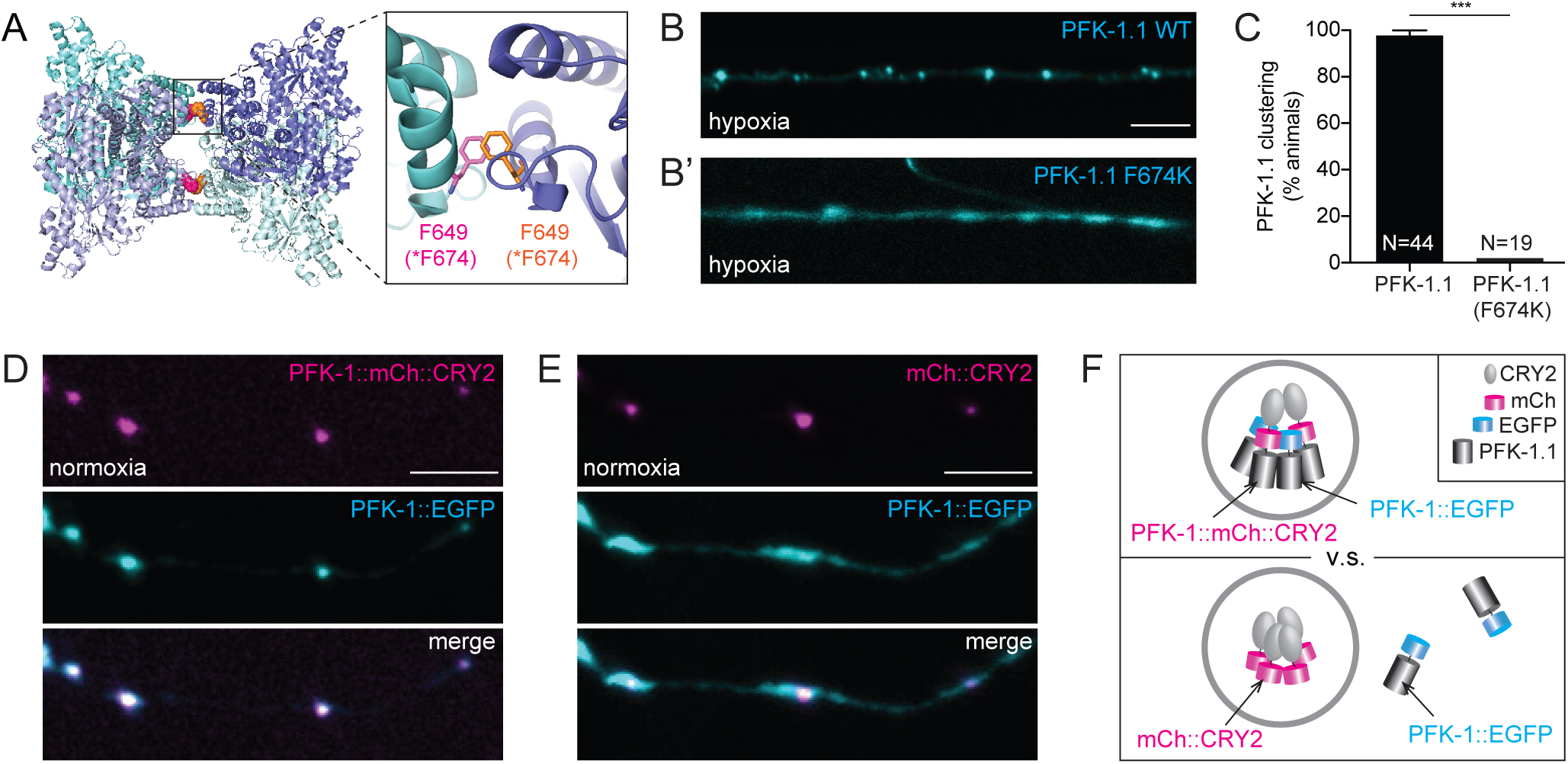
Multivalent interactions are necessary and sufficient for PFK-1.1 condensate formation. (**A**) Schematic of the tetramer structure of human PFK Platelet isoform (PDB code: 4XYJ) (Webb et al., 2015). Individual PFK subunits are colored using four different hues of blue. The F649 residue (equivalent to F674 in *C. elegans*), is shown in magenta and orange. One of the two sites of tetramer interface is zoomed in on the right to detail how F649 from each monomer interacts for tetramer formation. (**B, B’**) In **B**, PFK-1.1 condensates are observed after transient hypoxia, but, as shown in **B’**, PFK-1.1 F674K remains diffuse under the same conditions. (**C**) Percentage of animals that form PFK-1.1 condensates in response to transient hypoxia. An F674K mutation, predicted to affect PFK-1.1 tetramer formation, abrogates PFK-1.1 condensate formation. (**D**) Under normoxia, PFK-1.1::mCh::CRY2 (magenta, top panel) is sufficient to cause PFK-1.1::EGFP (cyan, middle panel) to form condensates. Merged image (bottom panel) shows co-localization of the CRY2-tagged (magenta) and non-tagged (cyan) PFK-1.1 condensates. (**E**) Under the same normoxic condition as in **D**, when co-expressed with mCh::CRY2 (magenta, top panel), PFK-1.1::EGFP (cyan, middle panel) remains diffuse. Merged image of the two is shown on the bottom panel. (**F**) Schematic of the localization of PFK-1.1::EGFP in the presence of PFK-1.1::mCh::CRY2 (top panel) or mCh::CRY2 (bottom panel). N = number of animals. All scales, 5µm. Error bars denote SEM. *, p < 0.05. **, p < 0.01. ***, p < 0.001 between indicated group.

To test whether the hydrophobic interface and multivalent interactions of PFK-1.1 are important for the formation of PFK-1.1 condensates, we mutated the *C. elegans pfk-1.1* F674 to a lysine residue (F674K). We first determined if *pfk-1.1* (F674K) allele was functional *in vivo* by generating a CRISPR allele and testing if this mutant affected, like other loss of function *pfk-1.1* alleles, the synaptic vesicle cycle (Jang et al., 2016). We observed that the allele was capable of sustaining the synaptic vesicle cycle under hypoxic conditions, suggesting that *pfk-1.1* (F674K) is functional *in vivo* (Figures S6B-S6D). Our findings are consistent with other *in vivo* studies demonstrating that PFK-1.1 tetramerization, while important for its biochemical activity *in vitro*, is not necessary for some of its activities *in vivo* (Marinho-Carvalho et al., 2006, 2009). Importantly, our study indicates that the *pfk-1.1* (F674K) allele makes functional protein that can be used to examine if tetramer formation is necessary for condensation.

Next, we examined the subcellular localization of PFK-1.1 (F674K)::EGFP. We observed that PFK-1.1 (F674K) proteins were incapable of forming condensates upon transient hypoxia, and remained diffusely localized (Figures 7B-7C and S6E). Our findings are consistent with tetramer formation being necessary for the oligomerization of PFK (Webb et al., 2017), and provide support for multivalent interactions being necessary for the formation of PFK-1.1 condensates.

To test if multivalent interactions are sufficient to induce the formation of PFK-1.1 condensates, we tagged PFK-1.1 with cryptochrome 2 (CRY2), a class of flavoproteins from *Arabidopsis thaliana* that are capable of mediating oligomerization (Bugaj et al., 2013). Optogenetic control of CRY2 has been recently used to drive phase separation of intrinsically disordered proteins (Shin et al., 2017). The extent of CRY2-induced oligomerization depends on the protein that the CRY2 is fused to, and it has been observed that tetrameric proteins highly enhance the homo-oligomerization of CRY2 (Che et al., 2015; Park et al., 2017). We observed that PFK-1.1::mCherry::CRY2 formed clusters even in the absence of any light stimulation and under normoxic conditions (Figure 7D). The PFK-1.1::mCherry::CRY2 clusters were functional and capable of rescuing the synaptic vesicle phenotype in *pfk-1.1(ola72)* mutant animals (Figure S6G). These results suggest that inducing PFK-1.1 functional self-association via CRY2 is sufficient to drive its condensation even in the absence of stimuli.

We then asked whether the PFK-1.1::mCherry::CRY2 ectopic condensates were sufficient to drive condensation of non-CRY2 tagged PFK-1.1. We examined this by simultaneously expressing PFK-1.1::mCherry::CRY2 and PFK-1.1::EGFP, or mCherry::CRY2 and PFK-1.1::EGFP, in single neurons. We observed that in neurons expressing mCherry::CRY2 and PFK-1.1::EGFP, PFK-.1.1 remained diffusely localized in the cytoplasm under normoxia, as expected (Figures 7E and 7F). Interestingly, in neurons co-expressing PFK-1.1::mCherry::CRY2 and PFK-1.1::EGFP, ectopic PFK-1.1::mCherry::CRY2 condensates contained PFK-1.1::EGFP even under normoxia (Figures 7D, 7F, and S6F). Our findings indicate that PFK-1.1 self-association, forced by the introduction of the CRY2 domain, likely drives a feed-forward loop that results in PFK-1.1 condensates even in the absence of stimuli. Together, our findings suggest that the multivalent interactions of PFK-1.1, mediated by hydrophobic domains that lead to its tetramerization, are the mechanisms by which PFK-1.1 phase separates *in vivo* into condensates.

## Discussion

PFK-1.1 localizes in subcellular compartments *in vivo*. Although widely regarded as cytosolic in nature, glycolytic enzymes like PFK have long been observed, both in biochemical and immunohistological studies, to self-associate into complexes and be enriched at subcellular compartments in specific cell types (Chu et al., 2012; Clarke and Masters, 1974; Green et al., 1965; Knull, 1978; Kurganov et al., 1985; Masters, 1984; Mercer and Dunham, 1981; Moses, 1978; Sullivan et al., 2003; Wilson, 1968, 1978). These observations led to the coining of the term “ambiquitous”, used to refer to the ability of glycolytic proteins to exist both in soluble and particulate/membrane-bound states (Wilson, 1978). Biochemical studies indicated that the state of the glycolytic proteins depended on the metabolic state (and identity) of the tissue from which they were harvested (Wilson, 1968, 1978). The ambiquitous properties of glycolytic proteins, however, remained controversial because the observed biochemical associations were weak, transient and dependent on the presence of particular metabolites (Brooks and Storey, 1991; Vas and Batke, 1981; Weber and Bernhard, 1982) and due to the lack of supporting *in vivo* evidence (Menard et al., 2014). In our study, and through a systematic examination of endogenous PFK-1.1 via the use of a hybrid microfluidic-hydrogel device, we conclusively determine that PFK-1.1 indeed displays distinct patterns of subcellular localization in specific tissues *in vivo*. Our findings are consistent with the hypothesis of the ambiquitous nature of glycolytic enzymes (Kurganov et al., 1985; Masters, 1984; Wilson, 1978), and with more recent cell biological studies which also observed compartmentalization of glycolytic proteins in response to specific stimuli (Araiza-Olivera et al., 2013; De Bock et al., 2013; Graham et al., 2007; Jin et al., 2017; Kohnhorst et al., 2017; Miura et al., 2013).

Our findings on the biophysical properties of PFK-1.1 during transient hypoxia, the fluid-like movements of the condensates, and the concentration dependence in their emergence indicate that PFK-1.1 clusters represent metastable liquid condensates. Interestingly, “aged” PFK-1.1 condensates slow down their dynamics, including non-fusing properties and decreased FRAP recovery. These findings suggest that changes in the metabolic state of the cell induced by hypoxia result in consistent (and predictable) changes in PFK-1.1 biophysical properties. Our findings also suggest that thermodynamic forces drive phase-separation during PFK-1.1 condensation. We note, however, that we also observed *in vivo* non-equilibrium properties to PFK-1.1 localization, which are likely the result of the interplay between cell biological regulatory mechanisms and phase separated condensates (what has been called “active emulsions” (Weber et al., 2019)). For example, we observed the exchange of material resulting, not in Ostwald ripening as would be predicted from thermodynamic equilibrium, but instead in similarly-sized adjacent condensates. We also observed that the localization of the PFK-1.1 condensates occurred near synapses, where PFK-1.1 was previously shown to be required for sustaining the synaptic vesicle cycle (Jang et al., 2016). These observations are consistent with a local “on-demand” assembly of PFK-1.1 condensates, similar to what has been described for other reaction-controlled assemblies of liquid compartments (Berry et al., 2018; Shin and Brangwynne, 2017). Importantly, our findings suggest that thermodynamic driving forces and regulatory biological mechanisms control the *ad hoc* formation of PFK-1.1 liquid condensates in neurites at specific subcellular compartments.

Multivalent interactions in PFK-1.1 mediate its condensation. Liquid-liquid phase separation can be induced via multivalent protein-protein and protein-RNA interactions (Alberti et al., 2019; Banani et al., 2017; Elbaum-Garfinkle et al., 2015; Langdon et al., 2018; Li et al., 2012; Nott et al., 2015; Smith et al., 2016; Wang et al., 2018; Zhang et al., 2015). PFK exists as a homotetramer capable of self-associating into higher-order structures (Webb et al., 2017). Similar to other self-associating proteins such as hemoglobin, PFK possesses unique geometrical symmetries that result in multivalent interactions that could contribute to phase separation (Chen et al., 2004; Galkin et al., 2002; Garcia-Seisdedos et al., 2017). Consistent with this hypothesis, we observed that mutations in PFK-1.1 in conserved sites predicted to disrupt tetramer formation and higher-order oligomers (Webb et al., 2015) abrogate PFK-1.1 capacity to form liquid condensates. Also consistent with this model, engineering a CRY2 self-association domain resulted in constitutive induction of PFK-1.1 condensates. Importantly, induction of PFK-1.1 condensates was sufficient to drive the localization of PFK-1.1 lacking the self-association CRY2 domain, suggesting that nucleation of PFK-1.1 condensates drive a feed-forward reaction that results in the compartmentalization of PFK-1.1 proteins into condensates. PFK tetramerization and self-association can be regulated by its substrate, by metabolites (including AMP and ATP), and by phosphorylation of its regulatory domain (Sola-Penna et al., 2010). ATP can also act like a biological hydrotrope to solubilize proteins (Patel et al., 2017). We hypothesize that a local change in metabolites (like AMP and ATP) could result in the local formation of PFK-1.1 condensates either by allosteric binding leading to PFK tetramerization or by the demonstrated hydrotrope capacity of ATP (or both). While we do not find RNA-interacting domains or intrinsically disordered regions in PFK-1.1, we note that glycolytic proteins have been found to associate with RNAs, and that this could also contribute to phase separation (Castello et al., 2016; Jin et al., 2017; Kastritis and Gavin, 2018; Mazurek et al., 1996). Together, our data are consistent with a model whereby PFK-1.1 regulated tetramerization in response to local metabolites might lead to multivalent interactions that drive its self-association into condensates.

PFK-1.1 condensates might represent a novel glycolytic compartment. PFK-1.1 does not co-localize with stress granule proteins nor does it form condensates under conditions known to cause stress granule formation. We had previously reported that PFK-1.1 co-localize with other glycolytic proteins, such as aldolase (ALDO) and glyceraldehyde 3-phosphate dehydrogenase (GAPDH) (Jang et al., 2016). In yeast, prolonged (24-hour) hypoxia results in PFK co-localizing with other glycolytic proteins into membraneless subcompartments termed G-bodies (Jin et al., 2017). The idea that glycolytic proteins compartmentalize into a glycolytic metabolon was first proposed after classical biochemical studies could not explain, based on known biochemical principles, the observed cellular rates of glycolysis (Moses, 1978; Sies, 1982). This led to the hypothesis that higher organizing principles, such as subcellular compartmentalization, must influence the observed flux for the glycolytic metabolic reaction in cells (Cori, 1956; Srere and Mosbach, 1974). While we show in this study that PFK-1.1 does not need to form condensates to drive the synaptic vesicle cycle, we hypothesize that the compartmentalization described here represents a mechanism that changes enzymatic parameters of the kinase to suit changing metabolic needs in the cell. Consistent with this, *in vitro* biochemical studies and modeling have demonstrated that the interaction between PFK and ALDO increases their enzymatic activities (Marcondes et al., 2011; Rais et al., 2000), and that higher order oligomerization of PFK is associated with an increase in its activity (Su and Storey, 1995). The subcellular organization and biophysical dynamics observed for PFK-1.1 might extend to other glycolytic proteins, and PFK-1.1 condensates might represent the hypothesized ‘glycolytic metabolon’, or a novel membraneless organelle that could regulate glycolysis.

Compartmentalization is a common way for cells to internally organize their reactions and cellular processes (Alberti, 2017; Schmitt and An, 2017; Walter and Brooks, 1995; Wombacher, 1983). Membraneless compartments in cells can serve multiple purposes, from inhibiting enzymatic activity by sequestering molecules to enhancing activity through mass action effects (Alberti et al., 2019; Banani et al., 2017; Case et al., 2019; Lee et al., 2013; Schmitt and An, 2017; Su et al., 2016). Although metabolic reactions by cytosolic proteins are generally thought of as distributed in cells, emerging evidence suggests that they could be compartmentalized in response to cellular states (Kastritis and Gavin, 2018; O’Connell et al., 2012; Prouteau and Loewith, 2018; Zecchin et al., 2015). For example, systematic studies on ∼800 metabolic enzymes in yeast identified widespread reorganization of proteins involved in intermediary metabolism upon nutrient starvation (Narayanaswamy et al., 2009). Enzymes involved in purine biogenesis are compartmentalized to enhance or inhibit their activity (An et al., 2008; Pedley and Benkovic, 2017). Asparagine synthetase paralogs and CTP synthase enzymes polymerize in response to cellular metabolic changes (Noree et al., 2014, 2019). We now show that PFK-1.1 dynamically compartmentalizes into condensates. The principles we uncover here on the subcellular organization of PFK-1.1 could represent important principles underlying the subcellular regulation of glycolysis, and conserved concepts on how metabolic processes are dynamically organized to modulate subcellular states.

## Acknowledgements

We thank Emmanuel Levy and Sucharita Dey (Weizmann Institute), Ceciel Jegers (Hyman lab), Robert Haase and Simon Alberti (MPI-CBG), Ann Cowan (UConn), Leslie Loew (UConn), John Kim and Mindy Clark (Johns Hopkins), Michael Rosen (UT Southwestern), Amy Gladfelter (UNC Chapel Hill), Joseph Hoffman (Yale University), Richard Goodman (Vollum Institute) and members of the Colón-Ramos lab for insightful discussions on the work and advice on the project.

We thank the Research Center for Minority Institutions program, the Marine Biological Laboratories (MBL), and the Instituto de Neurobiología de la Universidad de Puerto Rico for providing meeting and brainstorming platforms. Support for S.J. was provided by T32-GM007223. AA, AP and LJ were supported by a direct grant from the Max Planck Society. DRA and RL were supported by NSF EF1724026 and CBET1605679. Research in the DAC-R lab was supported by NIH R01NS076558, DP1NS111778 and by an HHMI Scholar Award.

## Movie Legends

**Movie 1. PFK-1.1 condensates induced by transient hypoxia reverts back to diffuse localization when the condition is returned to normoxia.** Transient hypoxia initiates at time 01:20 (minute:second), and also indicated by the appearance of the green square on the top right corner. Hypoxia is terminated 20 minutes later at 21:20. PFK-1.1 condensates can be seen dispersing after 21:20. See also Figure 3E. Scale, 5 µm.

**Movie 2**. **PFK-1.1 condensates repeatedly form and disperse during hypoxia-normoxia cycles.** Two rounds of 10 minutes of transient hypoxia (indicated by the appearance of a green square on the top right corner) interspaced with 5 minutes of normoxia are shown. A small region (white outlined box) of the neurite is zoomed in (2x; yellow outlined box) and the outline of PFK-1.1 localization in the same small region is shown on the bottom left corner. Elapsed time is shown in minute:second. See also Figure 3F. Scale, 5 µm.

**Movie 3. PFK-1.1 condensates undergo fusion.** The two PFK-1.1 condensates (in the white outlined box) can be seen fusing and relaxing into a spherical shape. A 2x zoomed inset is shown above. Elapsed time is shown in minute:second on the top left corner. Here, 00:00 is post 8 minutes of transient hypoxia. See also Figure 4C. Scale, 2 µm.

**Movie 4. PFK-1.1 condensates exchange material and undergo fusion.** The first PFK-1.1 condensate appear around 12 minutes into hypoxia (elapsed time shown as “minute:second” on the top left corner). Another one appears immediately to the right of it around 15 minutes post hypoxia, and the fluorescence intensity between the two condensates fluctuates until around 32 minutes post hypoxia, which then ends with a fusion event. See also Figure S5A. Scale, 2 µm.

**Movie 5. PFK-1.1 condensates exchange materials.** Four PFK-1.1 condensates appear in the white boxed region during transient-hypoxia as indicated by the elapsed time on the top left corner in minute:second. 2x zoomed in inset is shown immediately below. The condensates demonstrate fluctuation in fluorescence intensity, suggesting material exchange between the neighboring condensates. See also Figures 4E and 4F. Scale, 2 µm.

**Movie 6. PFK-1.1 condensates do not all fuse, but do have fluid-like movements.** Elapsed time is shown in minute:second on the top left corner. Here, 00:00 is post 30 minutes of prolonged hypoxia. PFK-1.1 condensates in the white boxed region, instead of fusing, “bounce” of each other. Note that compared to the fusion event (shown in Figure 4C and Movie3), which happened after approximately 10 minutes into transient hypoxia, the event recorded here is post 30 minutes of prolonged hypoxia. A 2x zoomed inset is shown at the bottom left. See also Figure S5C. Scale, 2 µm.

**Movie 7. PFK-1.1 clusters asynchronously appear in response to transient hypoxia.** Hypoxic condition, which is indicated by the appearance of a green square on the top right, starts at time 00:00 (minute:second). See also Figure 6A. Scale, 5 µm.

## Supplementary Figure Legends

**Figure S1 (related to Figure 1).**
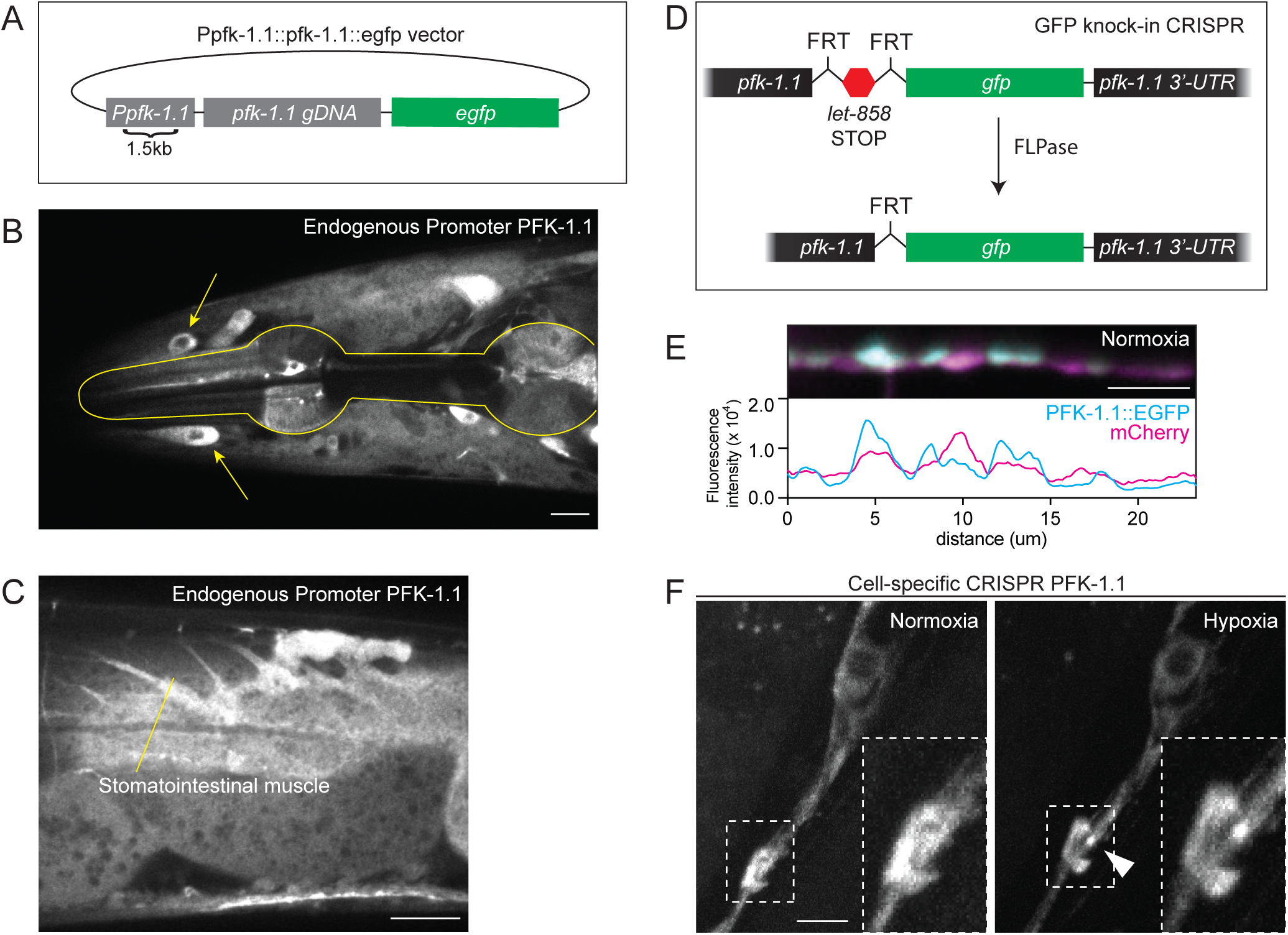
(**A**) Schematic of the expression vector for PFK-1.1:EGFP driven by its own promoter (Ppfk-1.1:PFK-1.1::EGFP). *Ppfk-1.1* contains 1.5 kilobase (kb) region upstream of the *pfk-1.1* start codon. *pfk-1.1 gDNA* is the genomic sequence of *pfk-1.1* that contains both introns and exons. (**B**) Head of *C. elegans* expressing the *Ppkf-1.1::pfk-1.1::egfp* array. Expression of PFK-1.1 can be seen in the pharyngeal muscle, as outlined, and in other unidentified cells (arrow). Scale, 10 µm. (**C**) Near the tail region of the animal, PFK-1.1 expression is observed in the stomatointestinal muscle. Scale, 15 µm. (**D**) Schematic of the strategy used for tissue-specific tagging of endogenous PFK-1.1 via conditional CRISPR. (**E**) Cytosolic mCherry (magenta) co-expressed with PFK-1.1 (cyan) in GABAergic neurons to observe the relative distribution of PFK-1.1. Note uneven enrichment of PFK-1.1 through different subcellular neuronal regions even in normoxic conditions. Line scan for PFK-1.1 and mCherry fluorescence level in lower panel. Scale, 5 µm. (**F**) The *unc-47* promoter was used in conditional CRISPR lines to tag the endogenous PFK-1.1 with GFP in a subset of tissues. After 30 minutes of hypoxia, PFK-1.1 clusters can be seen (arrowhead, right panel). Scale, 5 µm.

**Figure S2 (related to Figure 2).**
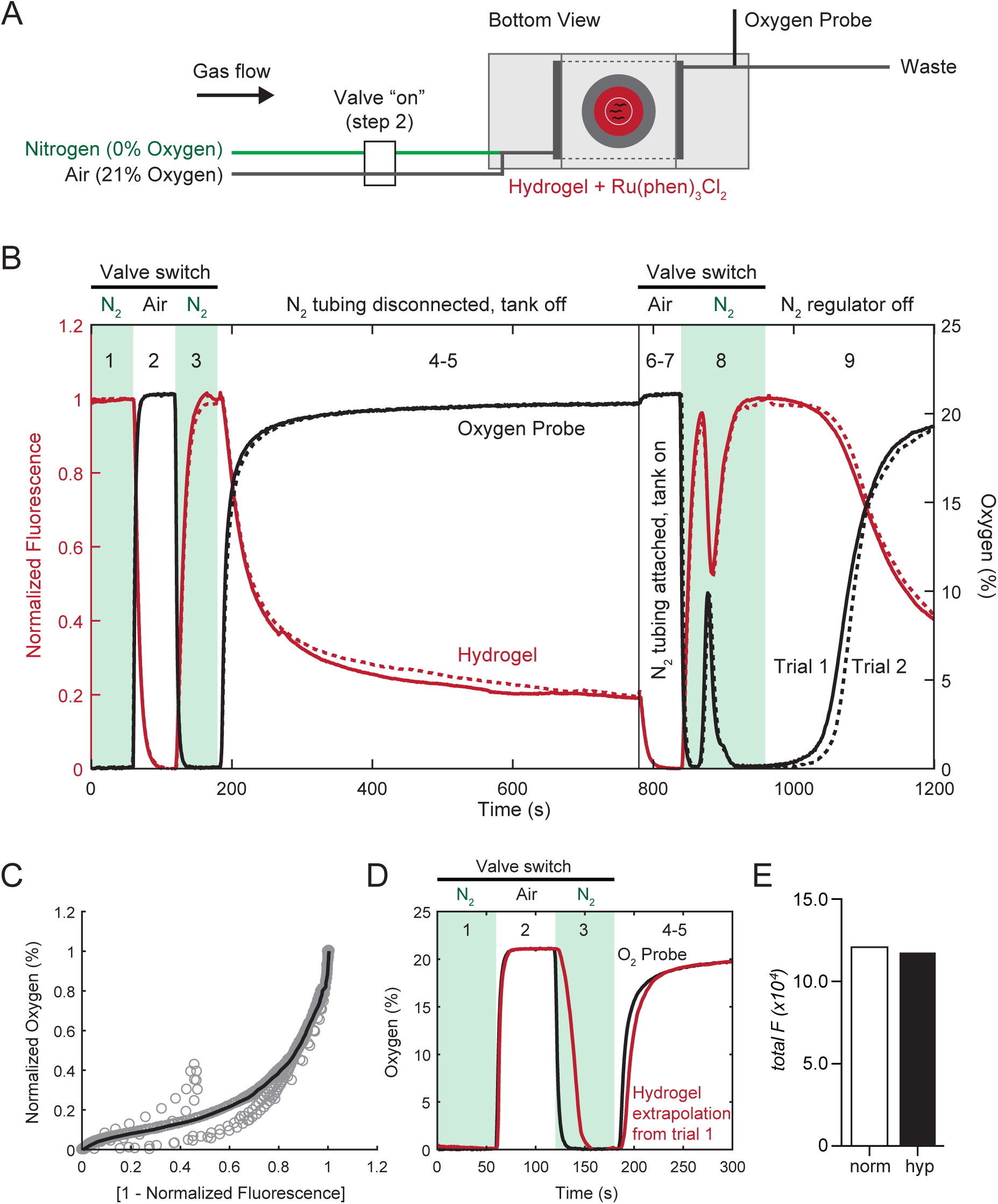
(**A**) Schematic of hybrid microfluidic-hydrogel device set up for calibration experiments. A pinch valve was used to control flow from compressed nitrogen and air tanks. When the valve was powered “on” using a custom Arduino microcontroller, air flowed through the chamber to the hydrogel and outlet (shown), in which oxygen concentration levels were continuously monitored using an oxygen probe. Synchronous changes in Ru(phen)_3_Cl_2_ fluorescence were monitored in the hydrogel by epifluorescence microscopy. When the valve was not powered, nitrogen flowed through the chamber (hypoxia) and outlet. (**B**) Line plot of synchronized oxygen probe and epifluorescence microscopy measurements (of Ru(phen)_3_Cl_2_ fluorescence) during a semi-automated sequence of valve switches. First, three valve-controlled switches from (1) nitrogen to (2) air to (3) nitrogen were delivered to observe steady-state minimum and maximum fluorescence intensities, followed by (4) manually turning off the nitrogen tank and (5) immediately disconnecting the nitrogen inlet tubing from the tank for 10 minutes. Next, a valve switch allowed (6) air flow to enter the device to determine shifts in baseline fluorescence. During this time, the nitrogen tubing was (7) re-attached and the tank was turned back on. Another valve switch (8) was used to restore low oxygen levels, yielding a short increase in oxygen likely due to air entering the previously disconnected nitrogen tubing. Finally, to compare differences in switching delays, the nitrogen regulator was (9) turned off and immediately back on after completing the 1200 second acquisition. The exact sequence was executed for a second trial (dashed line) and shown. (**C**) Scatter plot of all normalized oxygen concentrations larger than normalized fluorescence values versus [1 – normalized fluorescence]. Trend line represents the median values of binned oxygen concentration values, with linear interpolation between bins. (**D**) Estimated oxygen concentration in the hydrogel (using the calibration curve in **C**) compared with measured oxygen concentration in the gas channel for the first 300 s in **B**. (**E**) Total fluorescence of PFK-1.1 in Figure 2D neurite for normoxic condition (white box, “norm”) and transiently hypoxic conditions (black box, “hyp”). Note that there are no significant changes in the total levels of neurite fluorescence between the two conditions.

**Figure S3 (related to Figure 3).**
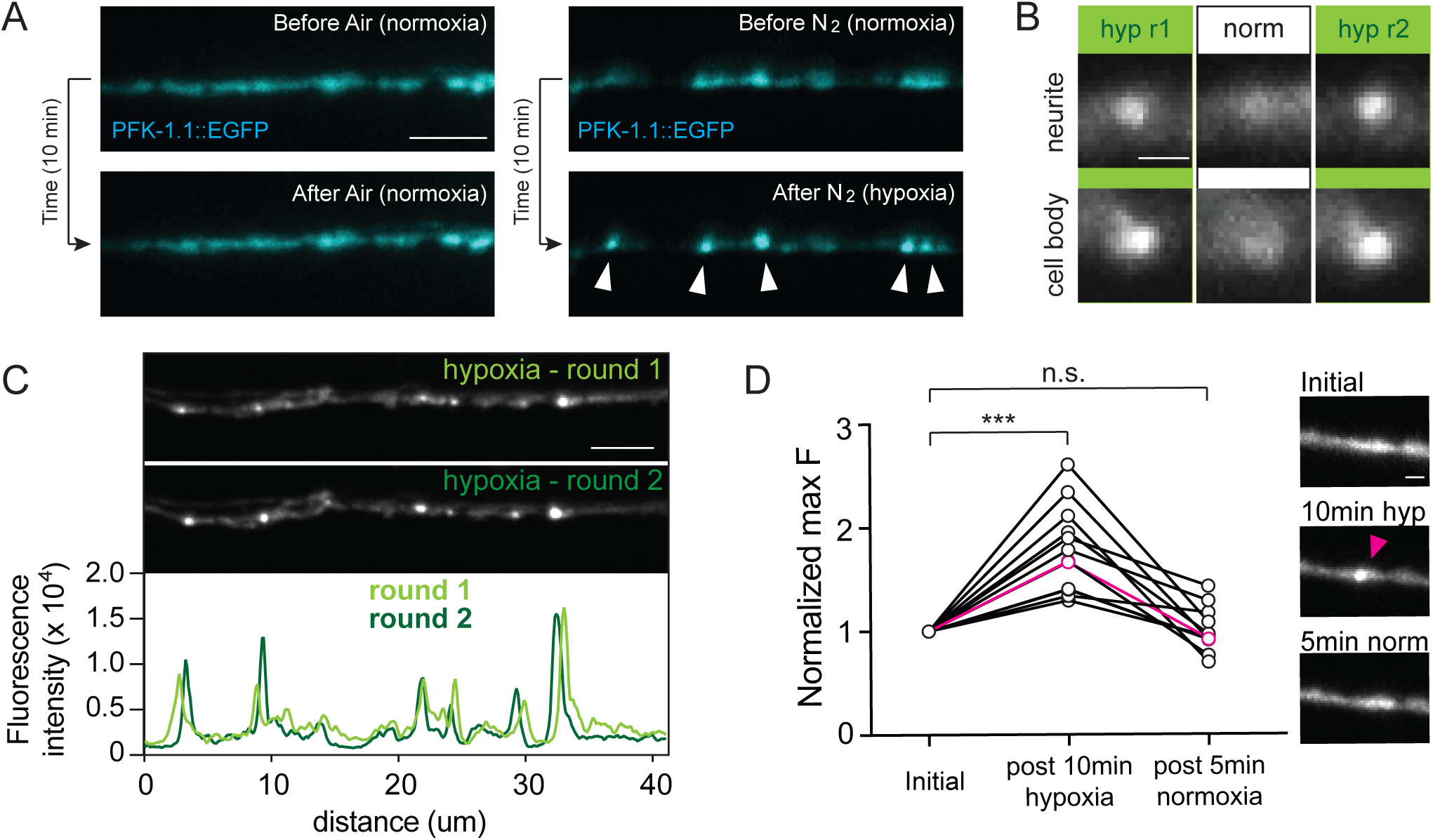
(**A**) PFK-1.1::EGFP before gas treatment (top panels) and after exposure to gas treatment for 10 minutes (bottom panels) with either normal air (left panels) or nitrogen gas (right panels). PFK-1.1 condensates (arrowheads) form specifically in response to transient hypoxic conditions. Scale, 5 µm. (**B**) PFK-1.1 condensates repeatedly appear in both the neurites (top panels) and cell bodies (bottom panels) of neurons through two rounds (r1 and r2) of normoxia (“norm”) and transient hypoxia (“hyp”) cycles. Scale, 1 µm. (**C**) PFK-1.1 localization after the first round of transient hypoxia (top panel) and the second round of transient hypoxia (middle panel). Note how PFK-1.1 condensates reappear at similar locations (this is another example of an additional neurite, as in Figure 2F and 2G). In lower panel, corresponding line scan for each round of transient hypoxia are shown: first round (light green) and second round (dark green). Scale, 5 µm. (**D**) Calculation of the normalized maximum fluorescence of twelve different neurite regions where PFK-1.1 condensates appear after 10 minutes of transient hypoxia and disperse after an additional five minutes of normoxia. Values were normalized to the initial maximum fluorescence value prior to transient hypoxia treatment. The normalized maximum fluorescence values showed significant increases by 1.79 +/- 0.12 fold (normalized mean fold +/- SEM; N=12 condensates) after transient hypoxia, and return to the basal level of 1.00 +/- 0.06 fold (normalized mean fold +/- SEM; N=12 condensates) after five minutes of normoxia. Right panels: image of the neurite prior to transient hypoxia treatment (top, labeled “Initial”), after condensate formation upon ten minutes of transient hypoxia (middle, labeled “10 min hyp”, with condensate pointed out by arrowhead), and upon five minutes of normoxia (lower, labeled “5 min norm”) corresponding to a representative condensate (in magenta) on the graph. Scale, 1 µm.

**Figure S4 (related to Figure 4).**
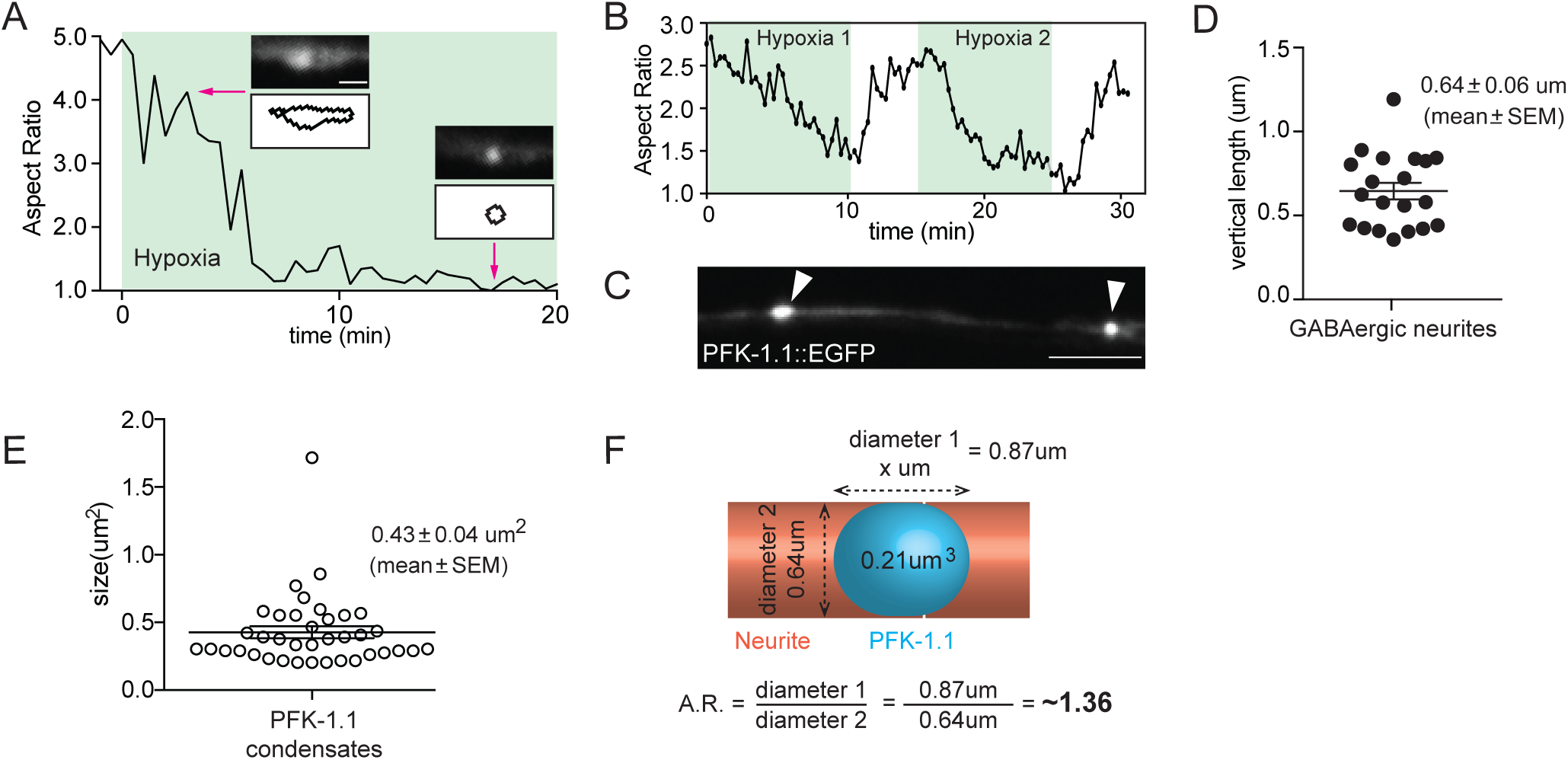
(**A**) Changes in the aspect ratio of PFK-1.1 for a given condensate over time. Once transient hypoxia was initiated (green region), the PFK-1.1 coalesced into a spherical structure. Two representative images (left: 3 minutes post-onset of transient hypoxia; right: 17.5 minutes post-onset of transient hypoxia) and corresponding outline of their morphology (white outlined boxes below) used to calculate the aspect ratio. Scale, 2 µm. (**B**) Changes in the aspect ratio of the PFK-1.1 condensates corresponding to Figure 3F over time. (**C**) Bigger PFK-1.1 condensates (left arrowhead) look like spherocylinders, as compared to smaller condensates (right arrowhead), which are spherical, suggesting that neurite space may constrain the shape of the fluid condensates. Scale, 5 µm. (**D**) We measured the diameter of GABAergic neurites, as visualized with cytoplasmic mCherry (also measured from electron micrographs, data not shown). Mean vertical length was 0.64 µm (N=20 neurite regions), which would only allow a perfect sphere with a volume of 0.14 µm^3^ to fit. (**E**) Measurement of PFK-1.1 condensate size shows an average area of 0.43 +/- 0.04 µm^2^ (mean +/- SEM; N=38 condensates) Assuming radial symmetry, its volume would be 0.21 µm^3^. (**F**) Schematic and calculation of aspect ratio for a theoretical spherical PFK-1.1 condensate with a volume of 0.21 µm^3^ that fits into a neurite with a diameter of 0.64 µm. Corresponding predicted aspect ratio is 1.36.

**Figure S5 (related to Figure 5).**
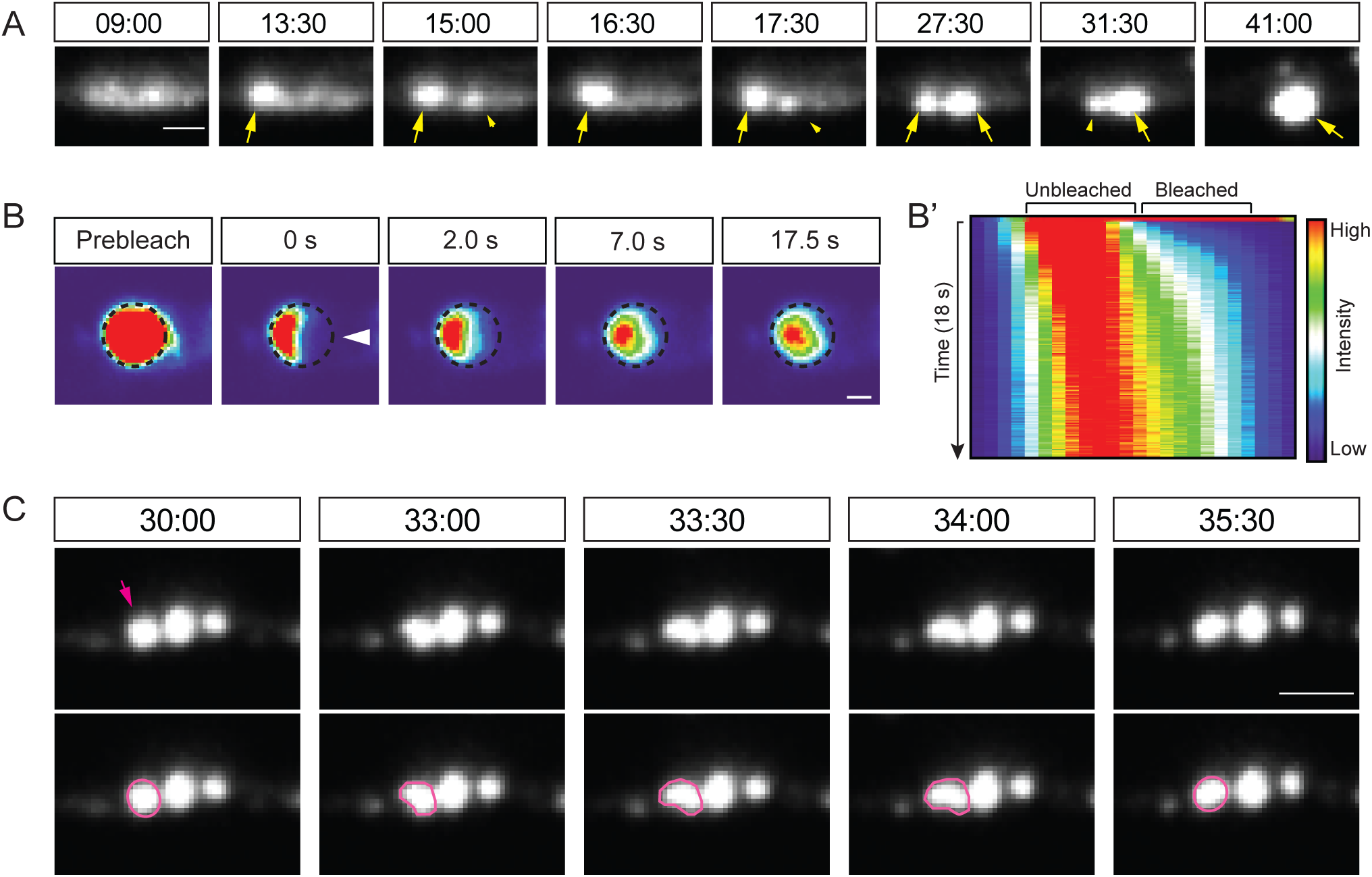
(**A**) Fluid dynamics of two neighboring PFK-1.1 condensates (yellow arrows) upon transient hypoxia (minutes:seconds after induction of transient hypoxia specified on top of the images). The first PFK-1.1 condensate appears in the second montage (13:30) and the second condensate appears soon after (third montage, 15:00). The fluorescence intensity between the two condensates fluctuates until around 32 minutes post hypoxia, which then ends with a fusion event between the two. See also Movie 5. Scale, 1 µm. (**B-B’**) Fluorescence recovery after photobleaching of a PFK-1.1 condensate in the cell body of a neuron also shows similar recovery dynamics to those observed in the neurites (compare with Figure 5A). PFK-1.1::EGFP condensate initial shape outlined with dashed circle, and partial area bleached highlighted with arrowhead in second panel. Scale, 1 µm. In **B’**, kymograph of the partially bleached condensate and recovery dynamics. (**C**) A PFK-1.1 condensate displays fluid motions (magenta arrowhead and outline in lower panels) as it bounces away from its closely neighboring condensate. Time elapsed since the start of transient hypoxia is indicated above in minutes:seconds. See also Movie 6. Scale, 2 µm.

**Figure S6 (related to Figure 7).**
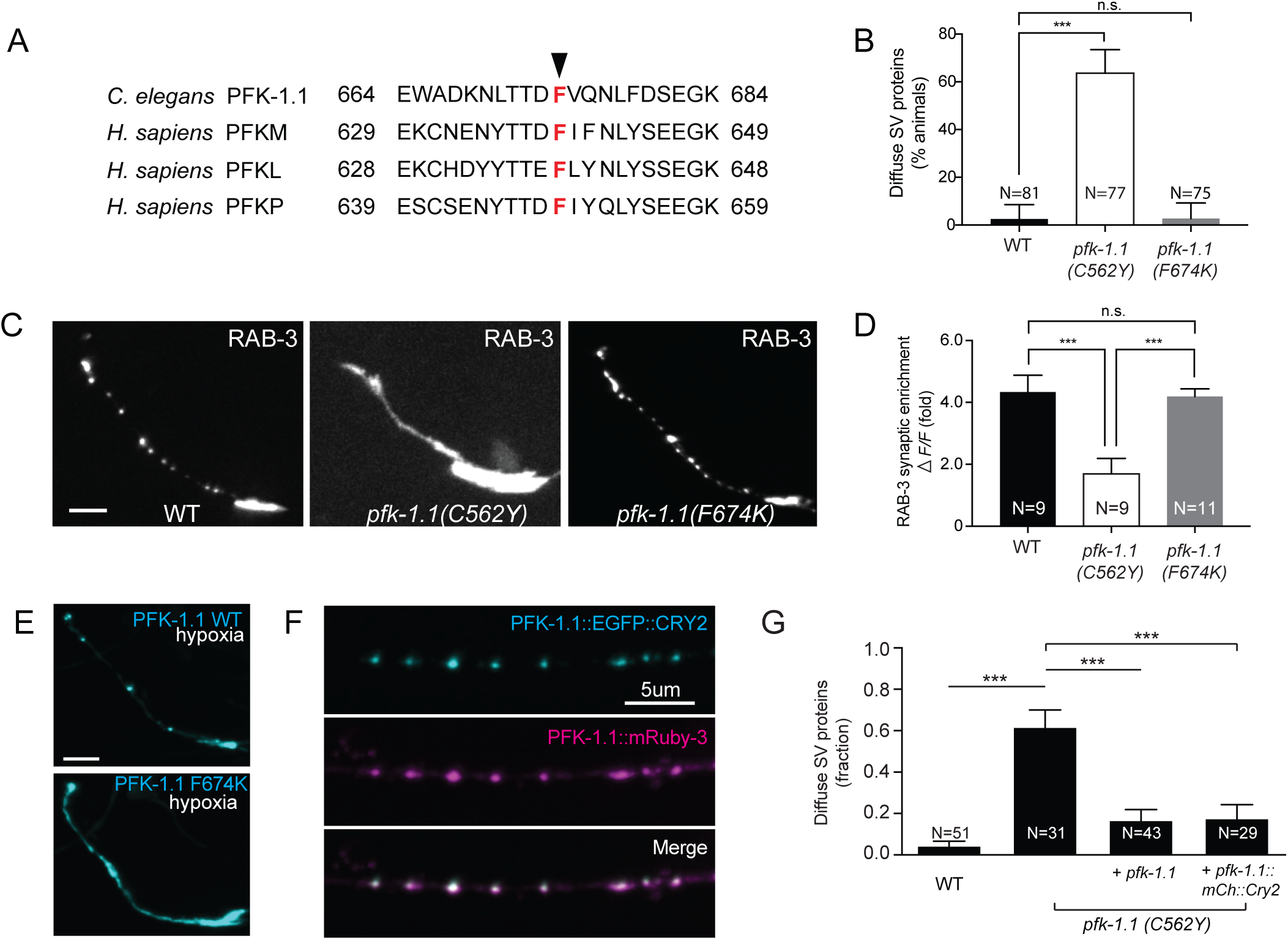
(**A**) Alignment of *C. elegans* PFK-1.1 with the three human isoforms of PFK, platelet (PFKP), muscle (PFKM), and liver (PFKL), in the region of the conserved F674 residue (red) necessary for PFK multivalent interactions and tetramer formation. (**B**) Percentage of animals displaying a diffuse pattern of RAB-3 in AIY neurons under hypoxia in wildtype (WT), *pfk-1.1* (C562Y) (also known as *pfk-1.1 (ola72)*) and *pfk-1.1* (F674K) mutants. As described before (Jang et al., 2016), PFK-1.1 loss of function mutants result in a diffuse RAB-3 pattern due to defects in synaptic vesicle endocytosis. Note how *pfk-1.1* (F674K) mutants do not phenocopy the loss of function *pfk-1.1 (ola72)* mutants for this phenotype, suggesting that tetramerization of PFK-1.1 is not necessary for this phenotype *in vivo*. (**C**) Images of the synaptic vesicle markers RAB-3 in AIY neurons of WT, *pfk-1.1* (C562Y), and *pfk-1.1* (F674K) mutant animals under transient hypoxia. RAB-3 displays a punctate pattern in WT and *pfk-1.1* (F674K) mutant animals, whereas it displays a diffuse pattern in the loss-of-function *pfk-1.1* (C562Y) mutant animals (which is the same as the allele *pfk-1.1(ola72)*). (**D**) Synaptic enrichment (**Δ**F/F; see Methods) of GFP::RAB-3 in AIY under hypoxia for the listed three genotypes. (**E**) WT PFK-1.1 in the AIY neuron forms condensates, whereas PFK-1.1 F674K in the AIY neuron remains diffuse under the same transient hypoxic conditions. (**F**) PFK-1.1::EGFP::CRY2 and PFK-1.1::mRuby-3 were co-expressed in GABAergic neurons. The top panel shows PFK-1.1::EGFP::CRY2 (cyan) puncta that formed under normoxia. In the middle panel, CRY2 untagged PFK-1.1::mRuby-3 also shows puncta (magenta). The bottom panel is a merged image of the two, demonstrating that they co-localize. (**G**) In transient hypoxic conditions, the percentage of animals displaying a diffuse pattern of RAB-3 in AIY neurons in WT, the loss-of-function allele *pfk-1.1 (C562Y)*, AIY cell-specific *pfk-1.1* array expressing *pfk-1.1(C562Y)*, and AIY cell-specific *pfk-1.1::mCh::CRY2* array expressing *pfk-1.1(C562Y)* animals. Note that the expression of *pfk-1.1* or *pfk-1.1::mCh::CRY2* both rescues the synaptic vesicle phenotype in *pfk-1.1(C562Y)* mutant animals, indicating that *pfk-1.1::mCh::CRY2* is functional and capable of rescue. N = number of animals. All scales, 5µm. Error bars denote SEM. *, p < 0.05. **, p < 0.01. ***, p < 0.001 between indicated groups.

## Materials and Methods

### *C. elegans* strains and transgenic lines

*C. elegans* were maintained at 20°C using OP50 *Escherichia coli* as a food source as previously described (Brenner, 1974). CRY2 expression clone carrying transgenic lines were kept separately in the dark in the same 20°C incubator. *C. elegans* N2 Bristol strain was used as wild-type.

For C-terminal endogenous tagging of PFK-1.1, a cell-specific CRISPR protocol (Dickinson et al., 2015; Schwartz and Jorgensen, 2016) was used to insert two flippase recombinase target (FRT) sites flanking *let-858* 3’-UTR that contains a transcriptional stop motif followed by a GFP sequence in front of the endogenous *pfk-1.1* 3’-UTR in the X chromosome (*pfk-1.1(ola368)*). Upon injection of cell-specific transgenes expressing FLPase, the *let-858* 3’-UTR containing the transcriptional stop motif is excised, leaving GFP fused to the C-terminus of PFK-1.1. To achieve specific expression of PFK-1.1::GFP in neurons and the body wall muscle, *unc-47* and *mig-13* promoters were used to drive the expression of FLPase in those cells.

For creating a point mutant of PFK-1.1, we used CRISPR strategies (Arribere et al., 2014; Paix et al., 2014) and generated *pfk-1.1 (ola379/F674K)* mutant animals. sgRNA sequence used was 5’ TCTGCACAAAATCTGTGGTG, and the repair template used was 5’ GCGAAACGCTACTTGG-TTGTCAGAAATGAGTGGGCCGACAAGAATCTCACCACAGATAAGGTGCAGAATCTTTTTGATTCTGAAGGAAAGAAGAACTTCACCACAAGAGTCAATG.

For generating expression clones, Gateway system (Invitrogen) and Gibson system (New England Biolabs) were primarily used. Transgenic strains (0.5–30 ng/μl) were made using standard techniques (Mello and Fire, 1995) and coinjected with markers *Punc-122::gfp, Punc-122::rfp, or Podr-1::rfp*. See Table 1 for all strains used in this study.

**Supplementary Table 1.** Strains used in this study.

| Strain Name | Genotype |
| --- | --- |
| DCR7312 | *olaex4409 [Ppfk-1.1:: pfk-1.1::egfp]* |
| DCR6732 | *pfk-1.1(ola368); olaex4038 [Pmig-13::FLPase]* |
| DCR6941 | *pfk-1.1(ola368); olaex4140 [Punc-47::FLPase]* |
| DCR6037 | *olaex3534 [Punc-47::pfk-1.1::egfp; Punc-47::snb-1::mCherry]* |
| DCR7155 | *olaex4296 [Punc-47::pfk-1.1::mRuby-3; Punc-47::tiar-1a::egfp]* |
| DCR7151 | *olaex4292 [Pttx-3::sl2::pfk-1.1::mRuby-3; Pttx-3::sl2::tiar-1a::egfp]* |
| DCR3921 | *olaex2271 [Punc-47::pfk-1.1::egfp; Punc-47::tom-20::mCherry]* |
| DCR6906 | *olaex4120 [Punc-47::pfk-1.1 F674K::egfp]* |
| DCR7038 | *olaex4219 [Pttx-3::sl2::pfk-1.1 F674K::egfp]* |
| DCR7799 | *olaex4731 [Pttx3::sl2::pfk-1.1::mCh::cry2; Pttx3::sl2::pfk-1.1::egfp]* |
| DCR7801 | *olaex4733 [Pttx3::sl2::mCh::cry2; Pttx3::sl2::pfk-1.1::egfp]* |
| DCR6444 | *olaex3817 [Punc-47::pfk-1.1::egfp::cry2; Punc-47::pfk-1.1::mRuby-3]* |
| DCR6426 | *olaex3802 [Punc-47::pfk-1.1::egfp; Punc-47::mCherry]* |
| TV1449 | *WyEx505 [Pttx-3::mCherry::erc; Pttx-3::gfp::rab-3]* |
| DCR1564 | *pfk-1.1 (ola72); WyEx505* |
| DCR7471 | *pfk-1.1 (ola379); WyEx505* |
| DCR6900 | *olaex4114 [Pttx-3::sl2::pfk-1.1]; pfk-1.1 (ola72); WyEx505* |
| DCR6987 | *olaex4180 [Pttx-3::gfp::rab-3; Pttx-3::pfk-1.1::mCherry::cry2]; pfk-1.1 (ola72)* |

### Hybrid microfluidic-hydrogel device set-up and calibration

A reusable microfluidic PDMS device was fabricated to deliver gases through a channel adjacent to immobilized animals, following protocols as previously described (Lagoy and Albrecht, 2015). A 50 µm, oxygen-permeable PDMS membrane was permanently bonded using air plasma to create an enclosed arena for gas flow, while the opposite side of the device was permanently bonded to a glass slide for structural integrity. Right-angle inlet and outlet holes were punched in the PDMS for ease of use with high-magnification inverted and upright microscopes. This reusable assembly was cleaned before each use by wiping the PDMS surface with ethanol, drying, and removing any remaining dust with tape. Animals were kept stationary during high-resolution imaging and exposure to shifting gas concentrations at the membrane surface by hydrogel immobilization (Burnett et al., 2018). To prepare the hydrogel assembly, a small volume (2 µL) of gel (20% PEGDA and 0.05% Irgacure 2959 in water) was pipetted onto the center of a hydrophobic glass slide containing a 100 µm thick PDMS spacer with a 6 mm diameter hole in the center, forming a rounded gel droplet. Glass slides or coverslips were rendered hydrophobic by 1 hour exposure to (tridecafluoro-1,1,2,2-tetrahydrooctyl) trichlorosilane vapor in a vacuum chamber, or “gel-adhesive” by 3 min exposure to 5% 3-(trimethoxysilyl)propyl methacrylate in ethanol (Burnett et al., 2018), which covalently grafts methacrylate groups to the glass that in turn covalently bind to the PEG chains of the hydrogel. Animals were then transferred into the solution and a gel-adhesive coverslip was placed over the hydrogel drop supported by the spacer. The assembly was placed over a UV light source (UVP, model UVGL-15, 4W) and illuminated for 2 minutes at 365 nm for gelation. The coverslip with spacer was carefully lifted off of the hydrophobic slide and a 2 µL drop of muscimol (50 mM in water) was then pipetted directly over the hydrogel disk. The gas device was carefully centered and lowered on top of the hydrogel assembly and clamped into a P2 series holder (Warner Instruments) to complete the closed system. Lastly, microfluidic tubing and components were connected to a nitrogen tank set to approximately 1-10 psi using a low-pressure regulator. To generate hypoxic conditions in the hydrogel, nitrogen flow was delivered continuously through the device assembly and confirmed by observing bubbles in a waste beaker filled with water connected with tubing to the device outlet. To generate normoxic conditions in the hydrogel, the nitrogen tank was turned off and the inlet tubing was immediately disconnected at the tank.

For device calibration, oxygen dynamics within the hydrogel were monitored by adding 4 µL of an oxygen sensitive fluorescent dye solution, 0.75 mM Ru(phen)_3_Cl_2_ dissolved in water, onto a 2 µL hydrogel disk before assembling with the gas device. Also, an oxygen probe (Ocean Optics HIOXY-PI600) was plugged into a second outlet to monitor corresponding oxygen concentration levels in the gas channel. A two-way solenoid pinch valve (NResearch 161P091) and custom Arduino controller were used to switch between constant flows of nitrogen or air (21% oxygen) through the device via separate inlets (Figure S2A). After this initial set-up, the oxygen probe was calibrated using a two-point calibration during nitrogen (0% oxygen) and air (21% oxygen) flow. MicroManager software was used to configure camera (2×2 binning and 1 second exposure) and acquisition settings (1 frame per second) for monitoring changes in fluorescence intensity through a 10X Leica objective (0.4 NA), RFP single-band filter set, detected by a Hammamatsu ORCA-ER mounted on a Leica DMI6000B microscope with EXFO X-Cite 120 Fluorescence Illumination System. A sequence of valve switches and tubing change steps were used to control shifts in oxygen concentration and monitored at 1 sample per second with the probe while synchronously recording changes in Ru(phen)_3_Cl_2_ hydrogel intensity (Figure S2B). This sequence was saved as a single TIFF stack file and repeated without disturbing the set-up and field of view for a second calibration trial. Mean fluorescence over the full field-of-view (672 × 512 px) centered on the hydrogel disk was measured for all 1200 frames and synchronized with oxygen concentration measurements from the outlet probe in the gas channel. Normalized fluorescence was calculated by scaling between the minimum fluorescence during steady 21% oxygen and the maximum fluorescence at steady 0% oxygen (as fluorescence scales inversely with oxygen). Slight increases in both minimum and maximum fluorescence of the oxygen sensor in the hydrogel were observed over time, likely due to a slow increase in concentration of the oxygen sensitive dye from evaporation. Scaling was adjusted by a linear interpolation of fluorescence envelope between the 0 – 21% oxygen transitions at minute 1 and near minute 15 (Figure S2B). To determine the calibration curve between hydrogel fluorescence and oxygen concentration, normalized fluorescence values were plotted against gas channel oxygen measurements, excluding data at times just following valve switches where hydrogel and gas channel concentrations would not be equal due to oxygen diffusion delays. The median fluorescence at binned oxygen concentration values (0.5% O_2_ bins) revealed a non-linear relationship between measured oxygen percent and fluorescence in the hydrogel (Figure S2C). This calibration curve was used to estimate oxygen concentration in the hydrogel based on linear interpolation between binned values (Figure S2D). From this, we estimate that switching between 21% and 0% oxygen in the hydrogel requires about 30 seconds to 1 minute, while switching between 0% and 21% oxygen in the hydrogel is nearly immediate. The directional difference in switching dynamics is likely due to the high oxygen concentration in PDMS (with high gas permeability) from ambient air, making it faster to increase oxygen, although hysteresis in dye fluorescence or kinetics could also contribute. All analysis was completed using MATLAB R2017a.

### Microscopy and image processing

Images of fluorescently tagged fusion proteins were captured live in *C. elegans* nematodes using a 60 CFI Plan Apo VC, numerical aperture 1.4, oil-immersion objective on an UltraView VoX spinning-disc confocal microscope (PerkinElmer Life and Analytical Sciences). Worms were immobilized using 50mM muscimol (Abcam) and hydrogel encapsulation (Burnett et al., 2018). ImageJ was used for image analysis, and it was used to adjust for brightness and contrast for image presentation. All the adjustments were kept identical for direct image comparisons unless otherwise stated. Maximum projections were used for all the confocal images, with the exception of the partial fluorescence recovery photobleaching (FRAP) experiments, which were in single planes. To correct for motion movement of the animal in time-lapse images, acquired image slices were mostly aligned using the Stack registration plugin (Thévenaz et al., 1998) in ImageJ or manually aligned and then the slices concatenated. Rendering of puncta was performed using Amira visualization software (Thermo Fisher Scientific), where gray values were used to create an isosurface superimposed on a gray-scale representation of the puncta in 3D (see Figure 4C). Finally, zoomed insets of images were obtained on ImageJ using the “Zoom in Images and Stacks” plugin developed by Gilles Carpentier.

### Quantification of phenotypic penetrance of glycolytic protein clustering and diffuse distribution of synaptic vesicle proteins

Animals were scored as displaying either “punctate” or “diffuse” phenotypes for the fluorophore-tagged glycolytic proteins or synaptic vesicle proteins after specified manipulations. Leica DM500B compound fluorescent microscope was used for the scoring. For hypoxia-induced conditions, coverslip-induced hypoxia was used as previously described (Jang et al., 2016) to calculate the percentage of animals displaying PFK-1.1 clusters or diffuse distribution of synaptic vesicle proteins. Statistical analyses were performed with Prism (GraphPad) and p values were calculated using Fisher’s exact test.

### Quantification of phenotypic expressivity of synaptic vesicle clustering

Quantification of synaptic vesicle clustering in Figure S6D was based on a previous protocol (Jang et al., 2016; Xuan et al., 2017). Briefly, fluorescence values for AIY neurites were obtained through segmented line scans using ImageJ. A sliding window of 2 mm was used to identify all the local fluorescence peak values and trough values. Synaptic enrichment was then calculated as % **Δ**F/F as previously described (Bai et al., 2010; Dittman and Kaplan, 2006).

### Fluorescence Recovery After Photobleaching (FRAP) of PFK-1.1

For FRAP, a 60 CFI Plan Apo VC, numerical aperture 1.4, oil-immersion objective on an UltraView Vox spinning-disc confocal microscope (PerkinElmer Life and Analytical Sciences) and Volocity FRAP Plugin were used. To calibrate the FRAP Plugin, steps laid out in Volocity User Manual (edition Sept 2011), “Calibrating the Photokinesis Accessory,” were followed.

For a full FRAP, z-stack images covering the entire volume of the neurite were acquired. A small, circular region, approximately the size of the PFK-1.1 punctum was used to bleach the entire PFK-1.1 punctum. Recovery images were acquired every 20 seconds. In ImageJ, acquired images were max projected. Images that passed the following three criteria were used for the analysis: 1) the fluorescence of the PFK-1.1 punctum was not saturated prior to photobleaching, 2) the punctum was photobleached to at least 80%, and 3) the neurite did not go out of focus during the image acquisition. For the fluorescence recovery calculations, a small ROI was drawn around the bleached punctum to measure its mean fluorescence. The mean fluorescence was then multiplied by the area of the ROI for total fluorescence. The same ROI was used to calculate a background fluorescence level. Background fluorescence was then subtracted from the total fluorescence for each time point. This calculated value was then normalized to the range of pre-bleach value as 1 and post-bleach value as 0 and plotted against time. In comparing the fluorescence recovery at different time points, *p values* were calculated using the Mann-Whitney *U* test.

For a partial FRAP, a small circular ROI was made and used for photobleaching: 17 (w, 2.05 µm) × 14 (h, 1.69 µm) x1 (d, 1 µm) dimension. For consistency, the same dimension was kept constant for all FRAP experiments. This ROI was placed just slightly overlapping with the PFK-1.1::EGFP punctum to cause partial bleach, but not full bleach of the entire punctum. Single plane images were acquired at maximum speed, resulting in approximately 0.1 second intervals for post-bleached images. Partial FRAP time-lapse images were analyzed in two ways: 1) kymograph to visualize the fluorescence recovery and 2) to individually calculate the fluorescence change in the bleached and unbleached regions. For calculating the fluorescence change, a small ROI was drawn within the bleached and unbleached region, and fluorescence was measured throughout time. Kymograph analysis was also performed using ImageJ.

### Viscosity estimation from partial FRAP

To understand the diffusion dynamics of PFK-1.1, we used partial photobleaching of PFK-1.1::EGFP in soluble state and in a condensate form. Because PFK-1.1 condensates were slightly bigger in the neuronal cell bodies, partial photobleaching and measurement of fluorescence recovery were conducted on PFK-1.1 found in the neuronal cell bodies. For normoxia condition, diffuse PFK-1.1 were photobleached and recovery measured. For transient hypoxia condition, PFK-1.1 condensates were induced with nitrogen gas for 10 to 20 minutes using the hybrid microfluidic-hydrogel device and then partially photobleached. From there, fluorescence recovery curves were obtained using the FRAP analysis plugin developed by Robert Haase for SCF-MPI-CBG. Using the fluorescence recovery, viscosity was approximated as previously described (Brangwynne et al., 2009). Briefly, the Stokes-Einstein relationship 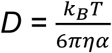 was used, where *D* is the diffusion coefficient, 𝑘_B_ is the Boltzmann’s constant, 𝑇 is the temperature, *α* is the radius of the PFK-1.1 particle (which we roughly estimated as 5 nm based on Webb et al., 2015), and 𝜂 is the viscosity. In comparing the calculated viscosity values, *p value* was calculated using the Mann-Whitney *U* test.

### Examination of subcellular localization of PFK-1.1 and stress granule protein TIAR-1

For heat shock induction, animals on NGM plates seeded with OP50 *Escherichia coli*, were incubated at 37°C for 1hour and were imaged immediately after that as previously described (Huelgas-Morales et al., 2016; Sun et al., 2011). For hypoxia induction, coverslip-induced hypoxia was used as previously described (Jang et al., 2016). AIY and GABAergic neurons were examined.

### Quantifications of subcellular localization of fluorophore-tagged proteins

For still images, the subcellular localization of fluorophore-tagged proteins, including PFK-1.1, were quantified using ImageJ and graphs plotted using Prism (GraphPad). 1) For the distribution of fluorophore-tagged proteins along the neurite, segmented line scans were performed and the graph of fluorescence intensity over distance was plotted. 2) In comparing the total amount of fluorophore-tagged proteins found in the neurite before and after hypoxia, an identical ROI around the neurite was drawn for those two time points. Total fluorescence values were obtained by multiplying the ROI with the mean fluorescence values found in the designated region. Finally, to account for any photobleaching, the raw total fluorescence value was normalized to the percent change of the background.

For time-lapse images, 1) a small ROI encapsulating individual area of the neurite where glycolytic clusters appear was designated in ImageJ and max fluorescence value, or pixel intensity, measured for each time point. The measured max fluorescence value was then plotted against time to show its change over time. 2) A segmented line was drawn through the neurite and kymograph was generated to show how PFK-1.1 protein localization changes under transient hypoxia using ImageJ. All the images used in the quantification analyses were obtained using identical microscopy settings.

### Aspect ratio calculations and size measurement of PFK-1.1

The aspect ratio (AR) was calculated with the max projected images. After thresholding the images to obtain an outline for each punctum, the *Analyze Particles* function in ImageJ was used to calculate AR values for each punctum and for all time points. Considering the resolution limit of a spinning disc confocal (approximately 300nm), any structure with a diameter less than 500nm and an area smaller than 0.2 µm^2^ was excluded from the analyses. PFK-1.1 condensate size or area was measured after thresholding the images and using the *Analyze Particles* function.

### PFK-1.1 rate of formation, dispersion and initiation time calculations

Using the acquired, time-lapse movies of PFK-1.1 cluster formation and dispersion with the hybrid microfluidic-hydrogel device, change in the max fluorescence of individual clusters were tracked over time using ImageJ. From plotting the max fluorescence change over time and performing linear regression, the region of where max fluorescence increases (for the formation) or decreases (for the dispersion) linearly was identified using the following parameters: minimum slope of at least 5 (fluorescence intensity / sec) and minimum length of half the time of the total treatment time. Any punctum with a starting fluorescence, or pixel, value of 1.2 × 10^5^ A.U. was excluded from the calculations because the max fluorescence did not further peak after this value or quickly reached a saturation point. MATLAB (MathWorks) was used to conduct linear regression with the criteria as mentioned above. Rate was defined as the slope of the fitted line and the initiation time was defined as the time for the first data point of the fitted line. The *p value* comparing the rate of PFK-11. cluster formation and dispersion was calculated using the Mann-Whitney *U* test.

### Normalization of max fluorescence

To compare how the max fluorescence changes after repetition of hypoxic and normoxic conditions, we normalized the max fluorescence of individual PFK-1.1 punctum at different time points. First, we identified the PFK-1.1 puncta that repeatedly appeared after the first and second round of hypoxic treatment (10 minutes of hypoxia). Then we measured the max fluorescence of individual punctum and the corresponding region for before hypoxia, after 10 minutes of hypoxia, and after an additional five minutes of normoxia. To account for photobleaching, we used a region that did not have PFK-1.1 puncta formation or localization change to normalize for each max fluorescence value. This value was then further normalized to the starting value or before hypoxic treatment to show the fold changes in max fluorescence through hypoxia and normoxia. *p values* comparing the max fluorescence at different time points were calculated using the Mann-Whitney *U* test.

### Neurite diameter measurement

To measure the diameter of the neurite, strains expressing cytosolic mCherry in GABAergic neurons were used. Total of 20 measurements was made by measuring the thickness of the fluorescent signal in the GABAergic neurites. Given the varicosity of the neurite, 10 total measurements were made in the synaptic bouton (mean average of 0.83 µm) and the other 10 measurements were made in the non-bouton areas (mean average of 0.45 µm). Total average was calculated by combining all 20 measurements. We also calculated the neurite diameter from GABAergic electron microscopy images in (Xuan et al., 2017).

